# Structural and functional analysis of the cerato-platanin-like effector protein Cpl1 suggests diverging functions in smut fungi

**DOI:** 10.1101/2022.07.21.500954

**Authors:** Paul Weiland, Felix Dempwolff, Wieland Steinchen, Sven-Andreas Freibert, Timo Glatter, Roman Martin, Gert Bange, Florian Altegoer

## Abstract

Plant pathogenic fungi are causative agents of the majority of plant diseases and can lead to severe crop loss in infected populations. Fungal colonization is achieved by combining different strategies, such as avoiding and counteracting the plant immune system and manipulating the host metabolome. Of major importance are effector proteins secreted by the fungi that fulfill diverse functions to support the infection process. Most of these proteins are highly specialized and structural and biochemical information is often absent. Here, we present the atomic structures of the cerato-platanin-like protein Cpl1 from *Ustilago maydis* and its homolog Uvi2 from *Ustilago hordei*. Both proteins adopt a double-Ψ-β-barrel architecture reminiscent of cerato-platanin proteins, a class so far not described in smut fungi. Our structure-function analysis shows that Cpl1 binds to soluble chitin fragments via two extended grooves at the dimer interface of the two monomer molecules. This carbohydrate-binding mode has not been observed previously and expands the repertoire of chitin-binding proteins. Cpl1 localizes to the cell wall of *U. maydis* and specifically enriches cell-wall degrading and -decorating proteins during maize infection. The architecture of Cpl1 harboring four surface exposed loop regions supports the idea that it might play a role in spatial coordination of these proteins. While deletion of *cpl1* has only mild effects on the virulence of *U. maydis*, a recent study showed that deletion of *uvi2* strongly impairs *U. hordei* virulence. Our structural comparison between Cpl1 and Uvi2 reveals sequence variations in the loop regions which might explain a diverging function.

## Introduction

Smut fungi constitute a large class of biotrophic plant pathogens that infect mostly grasses, among them many important cereal crops (Zuo et al., 2019). The individual smut fungal species have a narrow host range and establish a tight interaction for their parasitic lifestyle known as biotrophy (Benevenuto et al., 2018). *Ustilago maydis* infects the maize plant *Zea mays* and its ancestral form teosinte and forms dark black galls on infected parts of the plants that contain the mature teliospores (Brefort et al., 2009; Dean et al., 2012; Kämper et al., 2006). Plant infection by *U. maydis* is guided by the secretion of more than 400 effector proteins that allow fungal entry into the plant tissue, suppression of the plant immune system and metabolic rerouting for resource allocation (Lanver et al., 2017).

Effector proteins are generally classified as small, cysteine-rich proteins (typically between 200 and 300 amino acids) that harbor an N-terminal signal peptide for conventional secretion (De Wit et al., 2009; Lanver et al., 2017). Many of these proteins are host specific and lack conserved features. Furthermore, computational approaches to predict domains of known function or structure frequently fail to yield reliable results (Jones et al., 2018). In addition, effectors are often encoded in gene clusters or act in concert with other effectors, generating functional redundancy and single deletions might have little influence on the virulence of the deletion strains in infection experiments (Schirawski et al., 2010). The precise molecular functions of most effector proteins therefore remain enigmatic.

An important task of effector proteins from biotrophic fungi is the evasion of the plant immune system, especially during the early steps of host plant colonization. Microbial pathogens are recognized by host cell surface receptors through conserved microbe- or pathogen-associated molecular patterns (MAMPs or PAMPs) (Jones & Dangl, 2006). A prominent PAMP is chitin, an abundant component of the fungal cell wall (Pusztahelyi, 2018). Specialized receptors in the plant cell wall harboring LysM domains recognize chitin molecules which in turn triggers an immune response (Kaku et al., 2006; Kombrink et al., 2011) resulting in a broad range of cellular answers impairing fungal infections.

Consequently, fungal pathogens have evolved a repertoire of effector proteins that protect the fungal cell wall from plant chitinases or serve in scavenging of chitin-derived fragments. These effectors share binding properties towards oligosaccharides but are diverse in structure and function. The most prominent examples are proteins harboring LysM domains (Hu et al., 2021; Kombrink & Thomma, 2013), lectin-like domains (van den Burg et al., 2006) or proteins belonging to the cerato-platanin-like protein family (Pazzagli et al., 2014).

Cerato-platanins comprise a class of chitin-binding proteins exclusively found in filamentous fungi that exert functions in fungal development e.g., hyphal growth and cell wall remodeling (Chen et al., 2013; Pazzagli et al., 2014).

We here present the structural and molecular characterization of the **c**erato-**p**latanin-**l**ike protein 1 (Cpl1) from *U. maydis* and provide insights into its role during plant infection. Cpl1 is exclusively conserved among smut fungi and shows a high transcript abundance during the early stages of plant infection. Our study sheds light on the molecular details of Cpl1 and suggests that the cerato-platanin-like fold might be further distributed among fungi than anticipated so far. A structural comparison between Cpl1 and Uvi2 indicates that subtle changes in the sequence mapping to exposed surface loops might result in diverging functions of these effector proteins during maize and barley infection by *U. maydis* and *U. hordei*, respectively.

## Materials & Methods

### Accession numbers

The genes and encoding protein sequences are available from Uniprot (https://uniprot.org) und the following accession numbers: Cpl1 (A0A0D1E4Q7 [https://www.uniprot.org/uniprot/A0A0D1E4Q7]), Uvi2 (I2G262 [https://www.uniprot.org/uniprot/I2G262]).

### Strains and growth conditions

The *Escherichia coli* strain Dh5α (New England Biolabs) was used for cloning purposes. The *E. coli* strain SHuffle® (DE3) (Novagen) was used to express all produced proteins in this study. *E. coli* strains were grown under constant shaking in a temperature-controlled incubator. *Zea mays* cv. Early Golden Bantam (EGB, Urban Farmer, Westfield, IN, USA) was used for infection assays with *Ustilago maydis* (U.S.A., Ohio) (Kämper et al., 2006) and grown in a temperature-controlled greenhouse (light and dark cycles of 14 hours at 28 °C and 10 hours at 20 °C, respectively). *U. maydis* strains used in this study are listed in table S3. *U. maydis* was grown in YEPSlight medium (1 % (w/v) yeast extract, 0.4 % (w/v) peptone and 0.4 % (w/v) sucrose) at 28 °C with baffled flasks under constant shaking at 250 rpm or on potato dextrose-agar at 28 °C.

### DNA amplification and Molecular cloning

All primers and plasmids used in this study are listed in tables **S2** and **S3**. The open reading frames of all effectors cloned in this study were amplified from genomic DNA (*U. maydis* SG200). All genes were amplified without their predicted signal peptide (SignalP 5.0)(Almagro Armenteros et al., 2019) using specific primers (**Tab. S3**). A standard PCR protocol with Phusion® High Fidelity DNA polymerase (New England Biolabs) and primer-specific annealing temperatures was used for the DNA amplification. For the plasmid constructions, standard molecular cloning strategies and techniques were applied. Standard plasmid construction using restriction enzymes *BspH*I and *Xho*I (New England Biolabs) was used for *UMAG_01820*. Modular cloning using the restriction enzyme *Bsa*I and T4 ligase (New England Biolabs) was used to generate recombinant plasmids containing *UHOR_02700* and all other proteins used in this study. Briefly, vector and insert containing *Bsa*I recognition sites were digested with *Bsa*I at 37 °C for 4 minutes and ligated at 16 °C for 5 minutes. This reaction was repeated for 5 – 8 cycles with a final ligation step at 16 °C for 10 minutes. For plasmid amplification, recombinant plasmids were transformed in chemically competent *E. coli* DH5α strains (New England Biolabs). The correct sequence of all plasmids was verified using Sanger sequencing (Microsynth, Switzerland) with primers specified for the T7 promoter and terminator regions. For protein production *cpl1* was cloned into the pEMGB1 vector containing the solubility tag GB1, with a hexahistidine (6His) tag and a *Tobacco etch virus* protease (TEV) recognition site. *UHOR_02700* was inserted into pET24d for protein production.

### Transformation and generation of U. maydis strains

*U. maydis* protoplasts were transformed as previously described by Bösch *et al*. (2016). Briefly, 2 µg of donor DNA and 1 µg of plasmid in a volume of 10 µl ddH_2_O were added to *U. maydis* protoplasts and incubated on ice for 10 min. 500 µl of ice-cold sterile STC (1 M Sorbitol, 10 mM Tris-HCl pH 7.5, 100 mM CaCl_2_) solution supplemented with 40 % (v/v) PEG 3350 were added to the protoplasts and incubated for 15 min on ice. Transformants were plated on double-layered regeneration agar-plates. The first layer was supplemented with 4 µg/ml of carboxin topped with a layer of regeneration agar (10 g/l yeast extract, 20 g/l Bacto-pepton (Difco), 20 g/l sucrose, 1 M sorbitol, 15 g/l agar) without antibiotic. Cells were grown for 5 days and subsequently analyzed for successful transformation.

The gene *UMAG_01820* was disrupted in *U. maydis* SG200 using the CRISPR-Cas9 approach recently described for genetic manipulation of *U. maydis* (Schuster et al., 2016). A donor DNA was supplied during transformation to delete the respective open reading frame from the genome without further disrupting neighboring genes. Isolated *U. maydis* transformants were confirmed by colony PCR using the primers listed in **table S3** and sequencing (Microsynth, Switzerland). For the knockout generation, the plasmid pMS73 was digested with *Acc*65I to integrate the respective sgRNA expression cassette via Gibson Assembly, according to Schuster et al. (2016). The PCR obtained a double-stranded DNA fragment containing the respective target sequences, scaffold, terminator, and the corresponding overlapping sequences. The fragments were cloned into pMS73 (**Tab. S2**). The target sequences (**Tab. S3**) were designed using the E-CRISP tool (Heigwer et al., 2014).

The construction of HA-tagged *cpl1* was done as described for knockouts of *cpl1* but with a donor DNA encoding an HA tag with flanks designed for the C-terminus of *cpl1*. The inserts in all plasmids and knockouts were validated by sequencing.

### Plant infection assays

*U. maydis cpl1* KO-strains (SG200, FB1, and FB2) were grown in YEPS_light_ medium to an OD_600_ of 0.7 and subsequently adjusted to an OD_600_ of 1.0 using sterile double-distilled water. For the infection of maize plants, 500 µl of culture was injected into the stem of 7-day-old maize seedlings using a syringe as described by Kämper et al. (2006). In the case of FB1 and FB2, cultures of both strains were mixed 1:1 before injection. Disease symptoms of infected plants were scored at 12 days post infection (dpi) as described in (Kämper et al., 2006). Disease symptoms were quantified based on three biological replicates and are presented as stacked histograms.

### Immunolocalization

To localize Cpl1-HA in budding cells and filamentous hyphae, *U. maydis* strains expressing Cpl1-HA constitutively were suspended in 2% YEPSlight containing 0.1 mM 16-hydroxy hexadecanoic acid at a final OD_600_ of 0.5 and sprayed onto Parafilm. The Parafilm was placed on top of wetted paper towels inside square petri dishes and incubated at 28 °C for 17 h. The parafilm was washed with water and blocked with 1x phosphate-buffered saline (PBS) containing 3% (w/v) BSA and then incubated in α-HA antibody (Sigma; 1:1.500 dilution) diluted in PBS buffer and 3% (w/v) BSA at 4 ° C overnight. The samples were washed with PBS buffer and incubated in the goat anti-mouse IgG secondary antibody conjugated with Alexa Fluor 488 (Life Technologies; 1:1.500 dilution) for 1 h at 4 °C. After washing, the samples were analyzed using a Leica SP8 LSM confocal microscope equipped with a 100X objective (NA 1.4). Fluorophores were excited with a pulsed white light laser source at 488 nm. Photon emission was detected with a hybrid detector at the appropriate wavelength (495-530 nm). Images were processed with the Leica LAS AF software.

### Fungal stress assays

Fungal stress assays were performed as described previously (Weiland & Altegoer, 2021). Briefly, fungal strains were grown in YEPSlight medium [1% (w/v) yeast extract, 0.4% (w/v) peptone, and 0.4% (w/v) sucrose] to an OD600 of 1.0. The cells were pelleted and resuspended in sterile double-distilled H2O to an OD600 0.1. For the stress assays, 5 µl of the culture and indicated serial dilutions were spotted on CM (Holliday, 1974) plates supplemented with 50 µg/ml congo red, 45 µg/ml calcofluor white (Sigma-Aldrich), 1.5 mM hydrogen peroxide (H2O2), 1 M NaCl, or 1 M sorbitol. Images were taken after over-night incubation at 28°C.

### Protein production and purification

*E. coli* Shuffle (DE3) (Novagen) was transformed with pEMGB1-cpl1 to produce Cpl1 fused to an N-terminal GB1 tag including a hexahistidine tag. Transformed cells were grown on LB-agar plates (100 µg/ml ampicillin). Colonies from the plate were used as pre-culture in 100 ml LB medium (100 µg/ml ampicillin) and grown for 16 h at 37 °C under constant shaking at 180 rpm. The main culture was inoculated with the pre-culture to an OD_600_ of 0.1 and subsequently grown at 30 °C and 180 rpm to an OD_600_ of 0.6. The cultures were then cooled down to 20 °C and the protein production was induced by adding 0.5 mM isopropyl-β-D-1-thio-galactopyranoside (IPTG). The cells continued to grow for 20 h at 20 °C and 180 rpm. The cultures were harvested by centrifugation (4,000 *g*, 15 min, 4°C), resuspended in Buffer A (20 mM HEPES pH 8, 20 mM KCl, 40 mM imidazole and 250 mM NaCl) and subsequently disrupted using a microfluidizer (M110-L, Microfluidics). The cell debris was removed by centrifugation (50,000 *g*, 20 min, 4 °C). The supernatant was loaded onto Ni-NTA FF-HisTrap columns (GE Healthcare) for affinity purification via the hexahistidine tag. The columns were washed with Buffer A (10x column volume) and eluted with Buffer B (20 mM HEPES pH 8, 20 mM KCl, 250 mM imidazole and 250 mM NaCl). Prior to size-exclusion chromatography (SEC), the GB1-tag was cleaved off by adding 0.4 mg purified TEV directly to the eluate and incubating under constant rotation at 20 °C for 3 hours. Cleaved His-tagged GB1 and remaining TEV were removed via a second Ni-NTA purification after buffer exchange to Buffer A using an Amicon Ultra-10K centrifugal filter (Merck Millipore). The tag-free protein was subjected to SEC using a Superdex S200 Increase 26/600 column equilibrated in HEPES buffer (20 mM HEPES pH 7.5, 20 mM KCl and 200 mM NaCl). The peak fractions were analyzed using a standard SDS-PAGE protocol, pooled, and concentrated with Amicon Ultra-10K centrifugal filters.

### Protein crystallization

Crystallization was performed using the sitting-drop method at 20 °C in 0.5 – 0.75 μl drops. The crystallization drops contained the protein and precipitant solutions in either 1:1 or 1:2 ratio. Crystallization drops were set automatically using the Crystal Gryphon robot from Art Robbins Instruments. NeXtal JCSG suites I – IV and Classics II were used to screen for crystallization conditions. Native Cpl1 crystallized at 0.7 mM concentration within 21 days in 0.8 M LiCl, 0.1 M citrate pH 5.0 and 22,5 % (w/v) PEG 6000. Se-Met Cpl1 crystallized at 0.7 mM concentration within 1 month in 0.8 M LiCl, 0.1 M citrate pH 5.0, 20 % (w/v) PEG 6000 streak-seeded with native Cpl1 crystals. Uvi2 crystallized at 0.85 mM concentration within a week 0.1 M sodium acetate pH 5.0 and 10 % (v/v) MPD at a final pH of 5.0.

### Structure analysis by X-ray crystallography

Prior to data collection, the crystals were flash-frozen in liquid nitrogen employing a cryo-solution that consisted of crystallization solution supplemented with 15 % (v/v) glycerol. The data were collected under cryogenic conditions at the EMBL beamline P13 (Deutsches Elektronen Synchrotron; DESY). The data were integrated and scaled using XDS and merged with XSCALE (Kabsch, 2010). The structure of Cpl1 was determined by isomorphous replacement using data obtained from single-wavelength anomalous dispersion gathered by incorporating selenomethionine. The structure of Uvi2 was determined by molecular replacement in PHASER (McCoy et al., 2007) using the crystal structure of Cpl1 as a search model. Both structures were manually built in COOT (Emsley & Cowtan, 2004), and refined with PHENIX (Adams et al., 2010). The figures were prepared with PyMOL (Delano, 2002).

### Selenomethionine incorporation for anomalous diffraction

*E. coli* SHuffle T7 cells were transformed with pEMGB1-cpl1 grown on LB-agar plates infused with ampicillin (100 µg/ml) for 16 h at 37 °C. Colonies from the plate were used as a pre-culture of 400 ml LB medium (100 µg/ml ampicillin) grown for 16 h at 37 °C under constant shaking at 180 rpm. The cells were harvested at 4000 *g* for 15 min and resuspended in 10 ml M9 medium (37.25 g/l Na_2_HPO_4_, 16.5 g/l KH_2_PO_4_, 2.75 g/l NaCl, 5.5 g/l NH_4_Cl, pH 7.5). The resuspended cells were used to inoculate 5 l of M9 medium (100 µg/ml ampicillin) to an OD_600_ of 0.1. The M9 medium was infused with sterile and freshly made SolX solution (1 g/l L-lysine, 1 g/l L-threonine, 1 g/l L-phenylalanine, 0.5 g/l L-leucine, 0.5 g/l L-isoleucine, 0.5 g/l valine, 0.25 g/l selenomethionine, 80 g/l glucose, MgCl_2_, CaCl_2_). The cells were grown to an OD_600_ of 0.6 at 37 °C and 180 rpm. Protein production was induced by adding 1 mM IPTG. The cultures continued to grow at 37 °C and 180 rpm for 20 – 22 h. The cells were harvested by centrifugation and flash-frozen in liquid nitrogen to be stored at a temperature of -80 °C or immediately used for protein preparation.

### Molecular Docking

To virtually identify the putative binding pocket, molecular docking was carried out through AutoDock Vina (Trott & Olson, 2010). The receptor PDB file was prepared with AutoDockTools 4 (Morris et al., 2009) by adding the polar hydrogens and performing the conversion to PDBQT. Likewise, the ligand PDB file was also converted to PDBQT using AutoDockTools 4. The search grid box covered the whole receptor while the exhaustiveness parameter was set to 10000. Multiple AutoDock Vina runs with randomized seeds resulted in the same putative binding pocket again, indicating an informative prediction (Jaghoori et al., 2016).

### Analytical size-exclusion chromatography coupled to multi-angle light scattering (SEC-MALS)

SEC-MALS was performed using an Äkta PURE system (GE Healthcare) with a Superdex 200 Increase 10/300 column attached to a MALS detector 3609 (Postnova Analytics) and a refractive index detector 3150 (Postnova Analytics). The column was equilibrated with 0.2 µm filtered HEPES buffer for analysis at pH 7.5 or citrate buffer for studies at pH 5.0. The column was calibrated for apparent molecular weight determination using a mix of proteins with known molecular weights (conalbumin (75 kDa), ovalbumin (44 kDa), carbonic anhydrase (29 kDa), ribonuclease A1 (13.7 kDa), aprotinin (6.5 kDa)). The molecular weight was calculated by combining the refraction index and MALS values using a Debye fitting.

### Determination of dissociation constants by microscale thermophoresis (MST)

Dissociation constants of Cpl1 with different sugars were determined via microscale thermophoresis (MST). MST experiments were performed in HEPES buffer containing 0.06 % (v/v) tween 20 using a Monolith NT.115 with red LED power set to 100 % and infrared laser power set to 75 % (Jerabek-Willemsen et al., 2011). Tag-free Cpl1 was labeled according to the supplier’s instructions (dye NT 647, Nano-Temper Technologies). Subsequently, 500 nM of labeled protein was titrated against decreasing amounts of mannose, xylose, arabinose, chitobiose, chitotetraose, or cellobiose starting from 5 mM down to 0.15 µM. MST experiments were recorded at 680 nm and processed by NanoTemper analysis 1.2.009 and Origin8G.

### Hydrogen-deuterium-exchange mass spectrometry (HDX-MS)

Samples for HDX-MS were prepared by mixing 225 µl of purified Cpl1 (50 µM) with 25 µl of double-distilled water (apo state) or 25 µl of 50 mM concentrated chitobiose or chitotetraose, yielding a final ligand concentration of 5 mM.

Preparation of the HDX reactions was aided by a two-arm robotic autosampler (LEAP technologies). 7.5 μl of sample were mixed with 67.5 μl of D_2_O-containing SEC buffer (20 mM HEPES-Na pH 7.5, 20 mM KCl, 20 mM MgCl_2_, 200 mM NaCl) to start the exchange reaction and incubated for 10, 30, 100, 1,000 or 10,000 at 25 °C. Subsequently, 55 µl of the reaction were withdrawn and mixed with an equal volume of quench buffer (400 mM KH_2_PO_4_/H_3_PO_4_, 2 M guanidine-HCl, pH 2.2) at 1 °C. 95 µl of the resulting mixture were injected into an ACQUITY UPLC M-Class System with HDX Technology (Waters) (Wales et al., 2008). Undeuterated samples were prepared by similar procedure (incubation for approximately 10 s) through 10- fold dilution of the protein samples with H_2_O-containing SEC buffer. The injected samples were flushed out of the loop (50 µl) with H_2_O + 0.1% (v/v) formic acid (100 µl/min) and guided to a protease column (2 mm x 2 cm) kept at 12 °C that was filled with a 1:1:1 mixture of the protease’s porcine pepsin, protease type XIII from *Aspergillus saitoi* and protease type XVIII from *Rhizopus sp*. immobilized to bead material. The resulting peptides were collected on a trap column (2 mm x 2 cm) filled with POROS 20 R2 material (Thermo Scientific) kept at 0.5 °C. After 3 min of digestion and trapping, the trap column was placed in line with an ACQUITY UPLC BEH C18 1.7 μm 1.0 × 100 mm column (Waters), and the peptides eluted at 0.5 °C using a gradient of H_2_O + 0.1% (v/v) formic acid (A) and acetonitrile + 0.1% (v/v) formic acid (B) at a flow rate of 60 μl/min as follows: 0-7 min/95-65% A, 7-8 min/65-15% A, 8-10 min/15% A, 10-11 min/5% A, 11-16 min/95% A. The eluting proteins were guided to a G2-Si HDMS mass spectrometer with ion mobility separation (Waters), and peptides ionized with an electrospray ionization source (250 °C capillary temperature, spray voltage 3.0 kV) and mass spectra acquired in positive ion mode over a range of 50 to 2000 m/z in HDMS^E^ or HDMS mode for undeuterated and deuterated samples, respectively (Geromanos et al., 2009; Li et al., 2009). [Glu1]-Fibrinopeptide B standard (Waters) was employed for lock-mass correction. During separation of the peptide mixtures on the ACQUITY UPLC BEH C18 column, the protease column was washed three times with 80 µl of wash solution (0.5 M guanidine hydrochloride in 4% (v/v) acetonitrile,) and blank injections performed between each sample to reduce peptide carry-over. All measurements were carried out in triplicate, i.e. separate HDX reactions.

Peptide identification and analysis of deuterium incorporation were carried out with ProteinLynx Global SERVER (PLGS, Waters) and DynamX 3.0 software (Waters) as described previously (Osorio-Valeriano et al., 2019).

### Co-immunoprecipitation and mass spectrometry

*U. maydis* strains FB1 and FB2 harboring *cpl1*-*HA* in its native locus were used to infect *Z. mays* plants. The control infection was done using *U. maydis* SG200 containing HA-tagged mCherry with the signal peptide and the promoter of *Um*Cmu1 (UMAG_05731) integrated into the *ip*-locus of SG200 (citation). Infected plant leaves were harvested 3 days post-infection and flash-frozen in liquid nitrogen. Frozen plant material was pulverized using a cryogenic mill (Retsch, MM400) with 50 ml beakers loaded with a metal sphere of 2 cm diameter. Pulverization took place for 1 min at 30 Hz and under cryogenic conditions. The plant powder was transferred to 50 ml falcon tubes and stored at -80 °C.

For the co-immunoprecipitation experiments (Co-IP), 2 g of plant powder were added to 6 ml of HNN buffer (50 mM HEPES pH 7.5, 150 mM NaCl, 50 mM NaF, 5 mM EDTA) freshly infused with 1 mM PMSF, diluted cOmplete ™ protease inhibitor cocktail from Roche (stock solution 1:100) and 1 % (w/v) polyvinyl-pyrrolidone K30. The solution was kept at room temperature. A Dounce-homogenizer (Carl Roth CXE1.1) was used to disrupt and dissolve the plant powder entirely before adding 0.1 % (v/v) NP-40, 0.5 % (w/v) deoxycholic acid sodium salt, 1 % (w/v) dodecyl-β-D-maltosid, 1 % (w/v) dodecyldimethylaminoxid, and a cell wall degrading enzyme mix (1 U of cellulase Onozuka-R10, Macerozyme R-10, and cellulase from *Aspergillus niger*). The solution rotated at room temperature for 30 min. Subsequently, the cell debris was spun down at 4200 *g* for 10 min at 4 °C. The supernatant was transferred to Eppendorf tubes and split into 1 ml aliquots, to which 15 μl of magnetic anti-HA beads (Pierce®, Thermo Scientific) were added and incubated while rotating at 4 °C for 30 min. The beads were separated from the lysate using a magnetic rack. The supernatant was discarded, and the beads were washed three times with 400 μl of HNN buffer (containing all ingredients mentioned above, except for the cell wall degrading enzymes). The beads were washed five times with 800 μl 100 mM ammonium bicarbonate and subsequently flash-frozen in liquid nitrogen for the subsequent analyses by mass spectrometry as described previously (https://pubmed.ncbi.nlm.nih.gov/33028835/). In short, purified proteins were digested on-beads using trypsin, followed by reduction (5 mM Tris(2-carboxyethyl)phosphin (TCEP)) and alkylation (10 mM iodoacetamide) of peptides. Peptides were further desalted by solid phase extraction on C18 reverse phase spin columns (Machery-Nagel) and subsequently analyzed by liquid chromatography-mass spectrometry (LC-MS). Peptides were first separated by an Ultimate 3000 RSLC nano and a Q-Exactive Plus mass spectrometer (both Thermo Scientific). Settings were set as described previously (https://pubmed.ncbi.nlm.nih.gov/30638812/). The gradient length was adopted. Peptides were separated over 40 min from 98 % solvent A (0.15% formic acid) and 2 % solvent B (99.85 acetonitrile, 0.15 % formic acid) to 35 % solvent B at a flow rate of 300 nl/min. Label-free quantification was carried out by MaxQuant (https://www.nature.com/articles/nprot.2016.136) using standard settings with variable (oxidized M, deamidated N,Q) and fixed modification (carbamidomethylated C). The resulting MaxQuant “proteinGroups.txt” output table was loaded into Perseus (v1.5.2.6) (https://www.nature.com/articles/nmeth.3901). For calculation of enrichment factors in samples versus controls, values for proteins not detected in the control were imputed using the imputation function from normal distribution implemented in Perseus in default settings (width, 0.3; down-shift, 1.8).

## Results

### Cpl1 exhibits characteristic features of an abundant U.maydis effector

Plant infection by smut fungi is guided by the secretion of many effector proteins (Lanver et al., 2017; Xia et al., 2020; Zuo et al., 2019). Genes encoding effector proteins are usually not expressed under non-virulent conditions but show a substantial increase in expression during infection (Kämper et al., 2006; Lanver et al., 2018). Some of the genes with the overall highest transcript abundance are peaking in expression in the early stages of plant colonization (1 – 2 days post-infection (dpi)), (**Fig. S1A**). In addition to some cell wall degrading enzymes (CWDE, e.g. Afu1 and Egl1; **Fig. S1A**) two genes with the highest transcript abundance at 2 dpi are *rsp3* and a core effector of unknown function encoded by the gene locus *UMAG_01820* which we termed *cpl1* (**Fig. S1A**). While the protein encoded by *rsp3* (repetitive-secreted protein 3) is an important virulence factor that decorates the fungal hyphae during infection and shields them from the activity of the maize antifungal proteins AFP1 and AFP2 (Ma et al., 2018), nothing is known on function or structure of the protein encoded by *cpl1*.

*Cpl1* (RefSeq: XP_011387768.1) encodes a protein of 240 amino acids with a predicted molecular weight (MW) of approximately 27 kDa. *In silico* analyses using the Consensus Constrained TOPology prediction webserver (CCTOP) (Dobson et al., 2015), and SignalP 5.0 predicted no transmembrane helices but a signal peptide (SP) of 21 amino acids in length at the N-terminus of the protein. The Basic Local Alignment Search Tool for proteins (BLASTp) (Li et al., 2015), identified homologs of *cpl1* only in related smut fungi belonging to the order of *Ustilaginales* and the class *Ustilaginomycetes* (**Fig. S1B**). These organisms are *Pseudozyma hubeiensis* SY62, *Sporisorium scitamineum, Sporisorium reilianum* SRZ2, *Sporisorium graminicola, Ustilago trichophora, Melanopsichium pennsylvanicum, Kalmanozyma brasiliensis* GHG001, *Ustilago hordei, Ustilago bromivora, Moesziomyces antarcticus, Moesziomyces aphidis* DSM 70725, and *Testicularia cyperi* with identities ranging from 72% to 43%, respectively (determined by CLUSTAL2.1) (Sievers et al., 2011). Notably, Cpl1 contains four conserved cysteines among all orthologs (**Fig. S1C**).

In a previous study Uvi2 (UHOR_02700) has been identified as essential for the virulence of *U. hordei* (Ökmen et al., 2018) but its molecular function remained unknown. Cpl1 and Uvi2 have a sequence similarity of 57.5%. Moreover, both genes are highly expressed 2- and 3-days post infection of *Zea mays* and *Hordeum vulgare*, respectively (Lanver et al., 2018; Ökmen et al., 2018) (**Fig. S1B**).

### The crystal structure of Cpl1 reveals a dimeric double-Ψ-β-barrel architecture

No structural information on proteins homologous to Cpl1 was available and computational approaches failed to identify structural motifs or domains of known function. Thus, we determined the crystal structure of Cpl1 at 1.8 Å resolution by X-ray crystallography employing selenomethionine single-wavelength anomalous dispersion (Se-SAD). The structural model of the protein could be built to completeness.

Cpl1 consists of four α-helices and eight β-strands (**Figs. 1A, B**) forming a double-Ψ-β-barrel (DPBB) surrounded by flexible loops. The two α-helices α1 and α2, and a β-hairpin consisting of β1 and β2, form the N-terminal part (NTD) of the protein (**Fig. 1B**). The β-hairpin is stabilized by a highly conserved disulfide bond established by Cys77 and Cys96 (**Fig. S2A**). The core of Cpl1 contains a β-barrel formed by the six β-strands β3 – β8, which is flanked by helices α3 and α4 (**Fig. 1B**). Six-stranded beta-barrels are found in several proteins. However, the involving parallel strands rarely form two Ψ-structures, known as a DPBB (Castillo et al., 1999). The first Ψ-structure consists of the loop connecting strands β3 and β4 and the strand β7, whereas the second Ψ-structure consists of the loop connecting strands β7 and β8 and the strand β3 (**Fig. 1B**). The second disulfide bond formed by Cys124 and Cys157 stabilizes the first Ψ-structure and the β5/β6 hairpin, which is also part of the β-barrel (**Fig. S2A**). Four loops protrude from the central structural elements termed loop regions (**Fig. 1B, LR1-4**).

**Figure 1.**
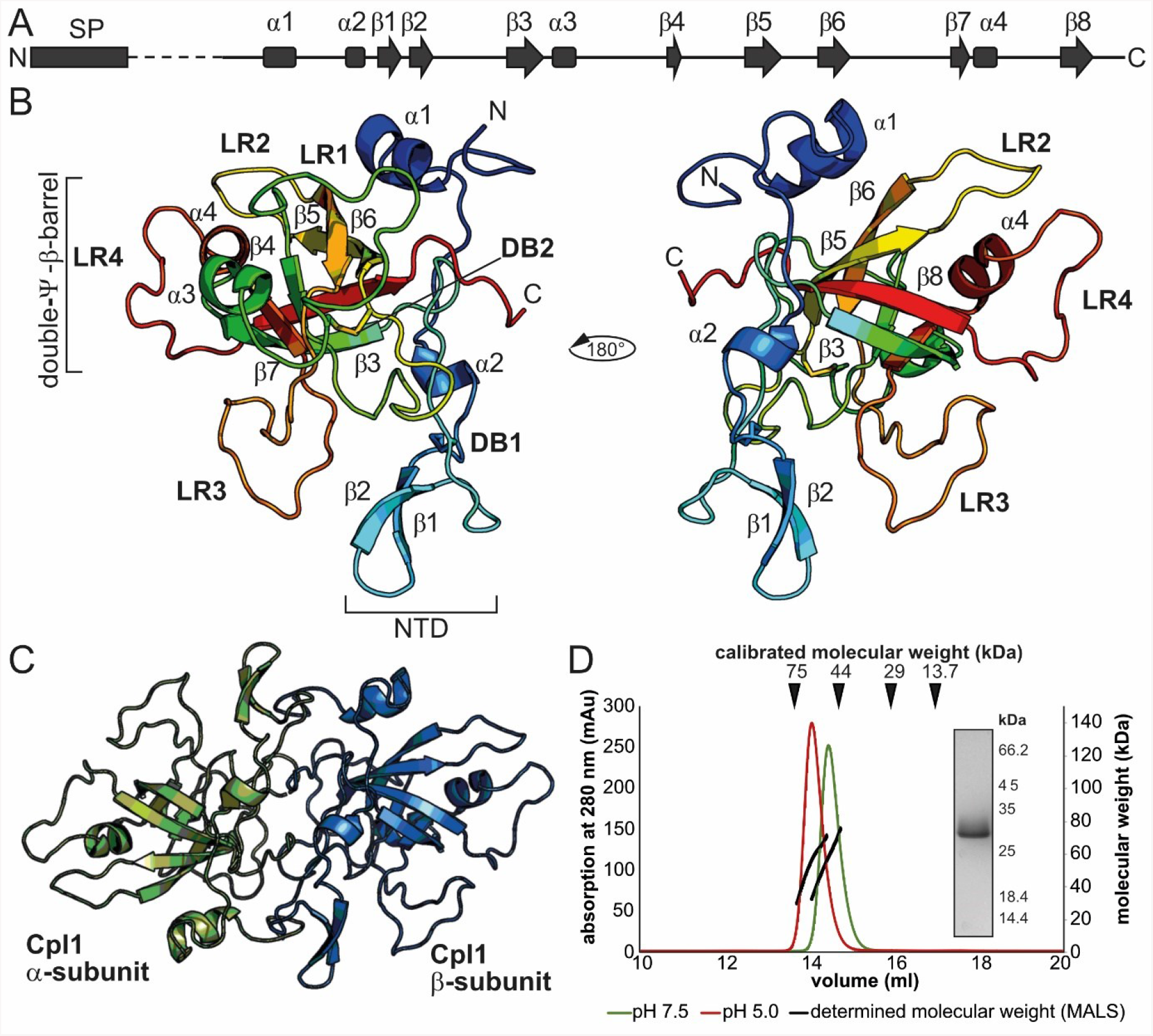
Crystal structure of Cpl1. **A**. Secondary structure of Cpl1. **B**. Cartoon model of Cpl1 colored in rainbow colors from N- to C-terminus. The two domains (NTD and DPBB) are indicated as well as the two disulfide bridges (DBs). Four loop regions (LR) extend from the central β-barrel. **C**. Cartoon model of the Cpl1 dimer with the two monomers colored in green and blue, respectively. **D**. SEC-coupled MALS analysis of Cpl1 at pH 7.5 and pH 5.0 shows the presence of a 50 kDa species. The inlet shows a Coomassie-stained SDS-PAGE of the peak fraction.

Notably, the asymmetric unit of the Cpl1 crystals contained six molecules that were suggested by PDBe PISA (Krissinel & Henrick, 2007) to form three dimers. The monomers of each dimer cover a buried surface area of 2000 Å^2^, implicating biological relevance (**Fig. 1C**). Four regions of Cpl1 contribute to the dimer interface between the two molecules. The first region covers residues 43 to 50 and partially aligns with residues 82 to 103 that form the second interface and residues 146 to 160 forming the third interface (**Fig. S2B**). Finally, the C-terminal residues 234 to 240 (region 4) are also buried in the dimer interface aligning mainly to residues of the second region (**Fig. S2C**). To assess whether Cpl1 forms stable dimers in solution, we employed size-exclusion chromatography coupled with multi-angle light scattering (SEC-MALS). SEC experiments were conducted at pH 7.5 and pH 5 to account for the rather acidic milieu of the apoplastic space. In both buffer systems, light scattering suggested masses of 53.9 and 54.5 kDa for pH 5.0 and 7.5, respectively. Our light scattering experiments thus reveal that Cpl1 forms a stable dimer at both pH values with the calculated molecular weight of a monomer being approx. 25 kDa (**Fig.1D**).

Taken together, our structural and biochemical analysis revealed that Cpl1 consists of a central DPBB surrounded by four loop regions stabilized by two conserved disulfide bonds. Two monomers form a stable homodimeric assembly via extensive interactions within two regions of the N-terminal domain and stretches in the C-terminal regions of the protein.

### Cpl1 has structural homology to ceratoplatanin-like proteins and binds to soluble chitin oligomers

With the structure of Cpl1 at hand, we set out to identify homologies to known structures that might allow to elucidate the precise function of this *U. maydis* protein. A search with the distance matrix alignment database (DALI) (Holm, 2020) retrieved a high structural similarity to several cerato-platatanin (CP) proteins from *Moniliophthora perniciosa*, a basidiomycete pathogen causing ‘witches’ broom disease of the cocoa tree (*Theobroma cacao*). More precisely, superposition to MpCp5 (PDB-Code: 3SUM) yielded a room mean square deviation (r.m.s.d) of 2.3 over 97 Cα-atoms (**Fig. 2A**). The superposition includes the six β-strands β3 – β8 as well as helix α4. Interestingly, despite some variations in the loop regions, the disulfide bond 2 (Cys124-Cys157) at Cpl1 superimposes almost perfectly with the first disulfide bond of MpCP5 (**Fig. S2D**). The N-terminus of Cpl1 containing the main dimer interaction interface does not superpose with parts of the MpCP5 structures. CPs are known to self-aggregate under specific conditions and some are also able to form dimers (Gaderer et al., 2014; Seidl et al., 2006). This might indicate a different mode of dimerization compared to Cpl1, where monomer formation was not observed.

**Figure 2.**
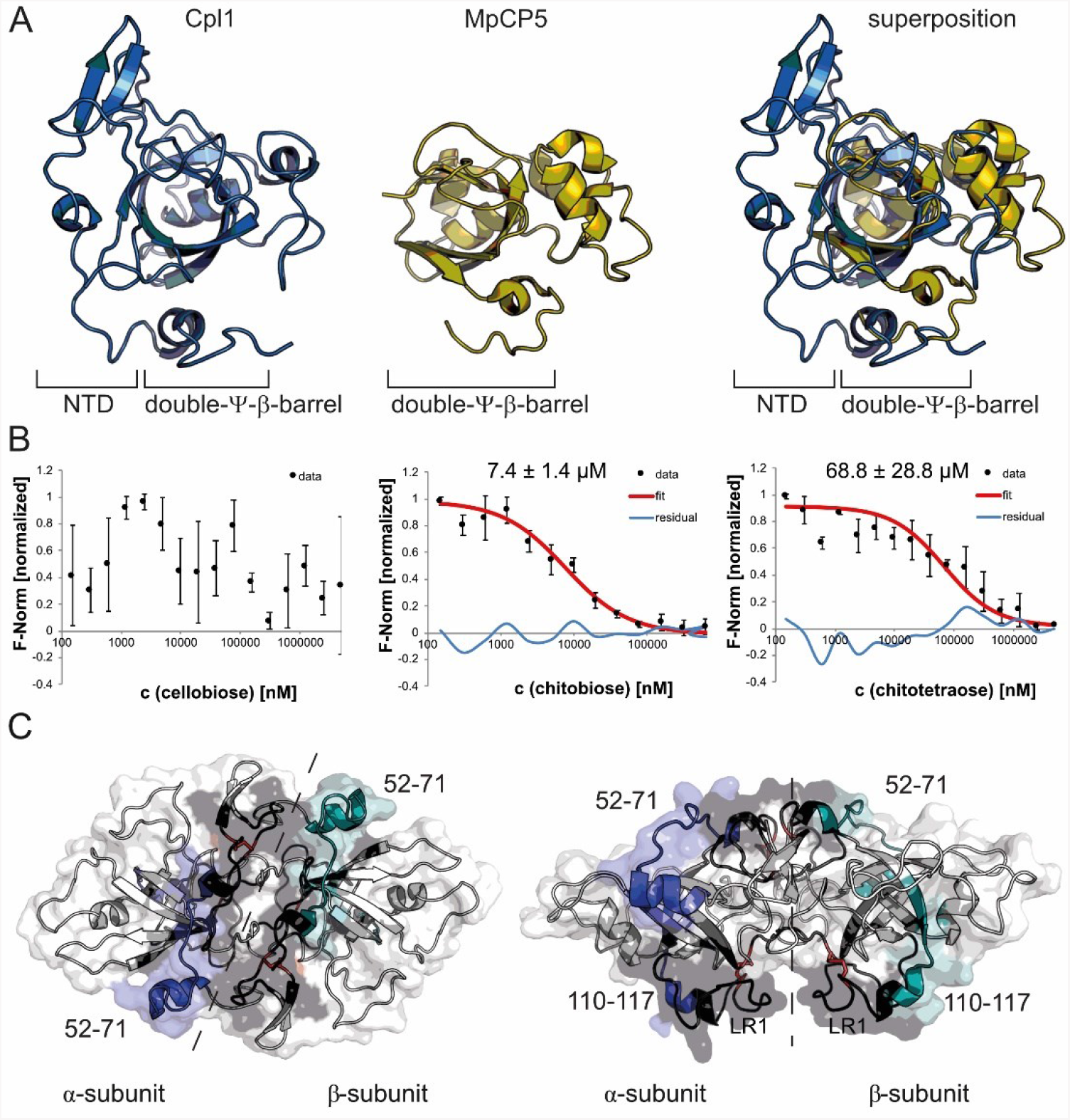
Cpl1 has high structural homology to cerato-platanin-like proteins and binds to chitin oligomers. A. Superposition of a Cpl1 monomer with MpCP5 (PDB-Code: 3SUM). B. Microscale-thermophoresis (MST) experiments with Cpl1. Shown are normalized fluorescence values (F-norm, black dots) of Cpl1 titrated against cellobiose, chitobiose and chitotetraose (concentration in nM). Where possible, a curve was fitted (red) used to calculate the dissociation constant (KD). Data points not included in the fitting are shown as residual data (curve identity, blue). The KD of Cpl1 binding chitobiose and chitotetraose were 7.4 µM (± 1.4), 68.8 µM (± 28.8), respectively. C. Areas of Cpl1 that exhibited reduced deuterium incorporation within at least two time-points in presence of chitotetraose (compare to Fig. S3) are colored in blue or deepteal for the α and β-subunits, respectively, and areas not covered by peptides in black. The side chains of the disulfide bridge-forming cysteine residues 77/96 and 124/157 are shown as red sticks.

CPs have been shown to interact with chitin polymers and its monomers specifically found in the fungal cell wall (Pazzagli et al., 2014). Chitin is a polymer of *N*-acetyl-D-glucosamin monomers linked by β-1,4-glycosidic bonds and based on the high structural homology towards CPs, we hypothesized that Cpl1 might also be able to bind chitin. Thus, we performed microscale-thermophoresis (MST) experiments to assess whether Cpl1 can bind to soluble chitin monomers, dimers and tetramers. Cpl1 was labeled with an amine-reactive dye and titrated with soluble chitin mono- and oligomer concentrations ranging from 5 mM to 152 nM. The estimated dissociation constant (K_D_) for the Cpl1-chitobiose interaction was 7.4 ± 1.4 µM and 68.8 ± 28.8 µM for the Cpl1-chitotetraose interaction, while no interaction of cellobiose with Cpl1 could be observed (**Fig. 2B)**.

To identify the binding interface of chitobiose and chitotetraose on Cpl1, we performed hydrogen-deuterium exchange coupled to mass spectrometry (HDX-MS). Specifically, the degree of deuterium incorporation of Cpl1 in presence of either ligand was compared to that of free Cpl1 and, after proteolytic cleavage of the protein into peptides, allowed for identification of the sites of ligand-dependent differences in HDX-MS (**Supplemental Dataset1-HDX**). In total, 79 peptides could be obtained that covered approximately 77% of the Cpl1 amino acid sequence (**Fig. S3A**). Very similar patterns of HDX reduction were apparent for chitobiose (**Fig. S3B**) and chitotetraose (**Fig. S3C**), in particular in peptides spanning residues 52-71 and 110-117 (**Fig. 2C, S4**). In the crystal structure of Cpl1 these residues locate to the upper and lower dimeric interface constituted by the two Cpl1 protomers (**Fig. 2C**). Unfortunately, peptides covering the regions surrounding the two disulfide bonds could not be retrieved due to incomplete proteolytic digest (**Fig. 2C**, black regions). This specifically reduced the information on binding events in the larger cleft involving LR1 (**Fig. 2C**, left panel). Our data thus suggest two regions within the Cpl1 dimer interface that are involved in binding of soluble chitin oligomers but further information was required to narrow down the specific binding location at Cpl1.

### A cleft in the dimer interface serves as major chitin binding interface

As suggested by our HDX-MS analysis, we had a closer look at the two clefts in the Cpl1 dimer interface. To substantiate our findings, we also performed molecular docking with chitobiose and chitotetraose using AutoDock (Trott & Olson, 2010). The different states of the docked molecules all located within the cleft formed by LR1 that extends from helix α3 to strand β3 (**Fig. 3A**). Combining our HDX-MS and molecular docking results, we chose amino acids that might be involved in the interaction to generate alanine variants and perform MST experiments. Specifically, we varied D69 and E72 into alanines that both locate within or close to helix α2 in the first binding interface identified by HDX-MS. Our MST experiments showed a ∼10-fold decrease in affinity towards chitobiose (66.31 ± 3.54 µM in Cpl1_D69A/E72A_ vs. 7.4 ± 1.4 µM in the WT) (**Fig. 3B**) while the affinity towards chitotetraose was seemingly not affected. However, when varying N151 and R155 to alanines as residues that reside in the prominent cleft on the opposite site of the first interface, we could not observe a K_D_ towards chitobiose anymore (**Fig. 3B**). Furthermore, the affinity towards chitotetraose was reduced by roughly 2-fold (110.54 ± 21.4 µM Cpl1_N151A/R155A_vs. 68 ± 28.8 µM in the WT). The two amino acids N151 and R155 are located on both ends of the cleft and the binding interface might therefore accommodate even larger chitin oligomers in the natural context (**Fig. 3C**). In conclusion, we can show that two regions within the interface between two Cpl1 protomers contribute to binding of soluble chitin oligomers, with the one formed by LR1 being the major one.

**Figure 3.**
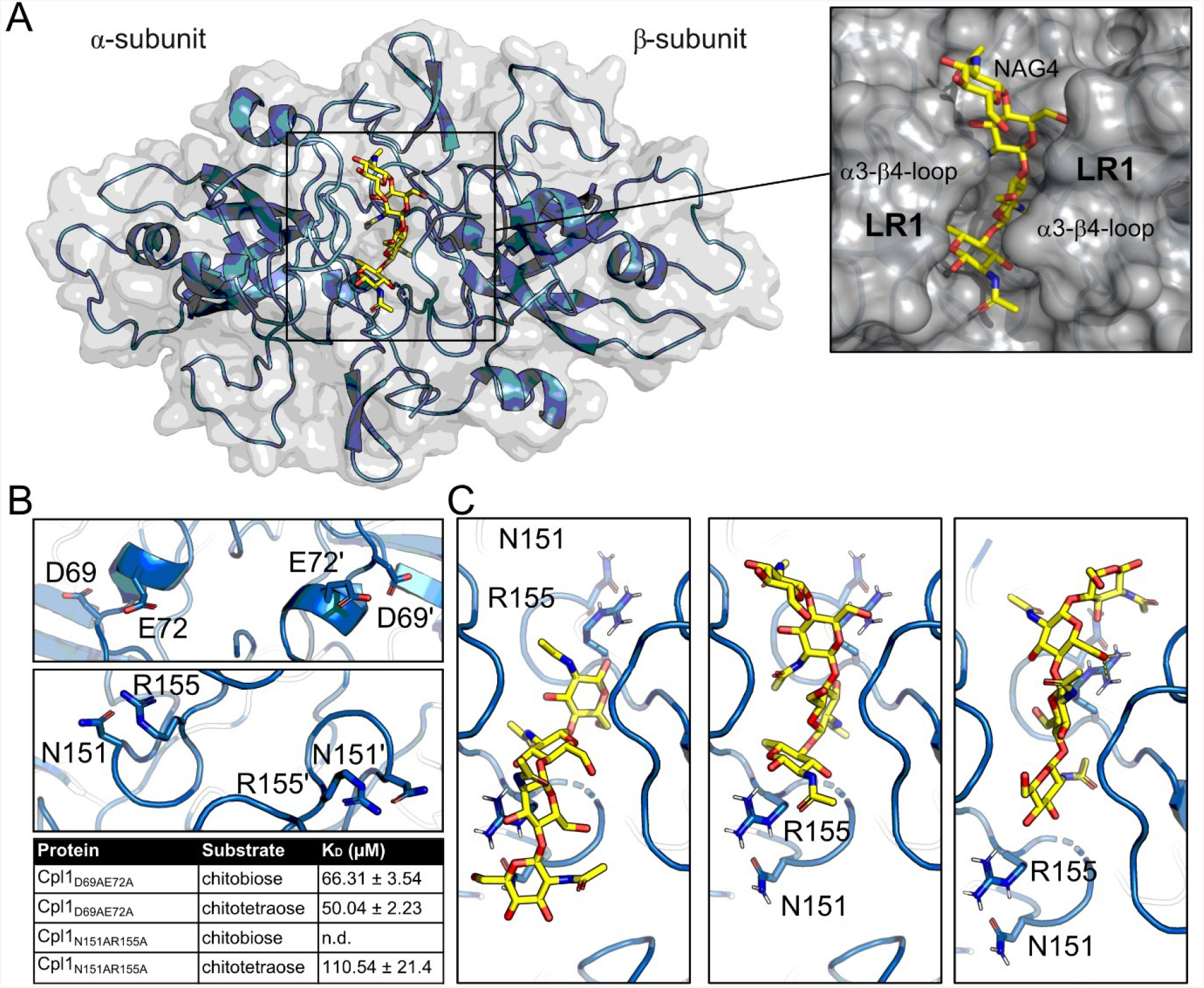
A groove in the interdimer interface at Cpl1 is important for binding of chitin oligomers. A. Cpl1 dimer (blue) with a chitotetraose docked into the interdimer groove (yellow). The inset shows a closeup of the chitotetraose squeezed between the α3-β4-loop (LR1) of both protomers. B. Amino acids that were varied to alanines and results of MST experiments employing Cpl1_N151A/R155A_ and Cpl1_D69A/E72A_. C. Closeup of differently docked chitotetraose molecules. N151 and R155 are highlighted and have been varied to alanines.

### Cpl1 is dispensable for sporidial growth of U. maydis and its deletion only mildly affects virulence

With the molecular details of Cpl1 at hand, we next aimed to understand how a deletion of *cpl1* would influence the growth of *U. maydis* and estimate the influence of Cpl1 on the virulence in plant infection experiments. Therefore, we first generated a *cpl1* deletion in the wildtype *U. maydis* FB1 and FB2 background (Banuett & Herskowitz, 1989) by using a CRISPR-Cas9-dependent approach (Schuster et al., 2016). We could not detect any phenotypic differences in disease symptoms between FB1xFB2 infected plants and plants infected with FB1Δcpl1xFB2Δcpl1. As the infection with FB1xFB2 leads to strong infection symptoms with a substantial number of dead plants, minor difference in virulence cannot readily be assessed. We therefore generated stains deleted for *cpl1* in the solopathogenic SG200 background. Indeed, we could detect subtle differences in the virulence of strains deleted for *cpl1* compared to the respective parental strains (**Fig. 4A**). While the number of plants with small tumors was significantly smaller in SG200Δcpl1 compared to SG200, the total number of larger tumors was higher (**Fig. 4A**).

**Figure 4.**
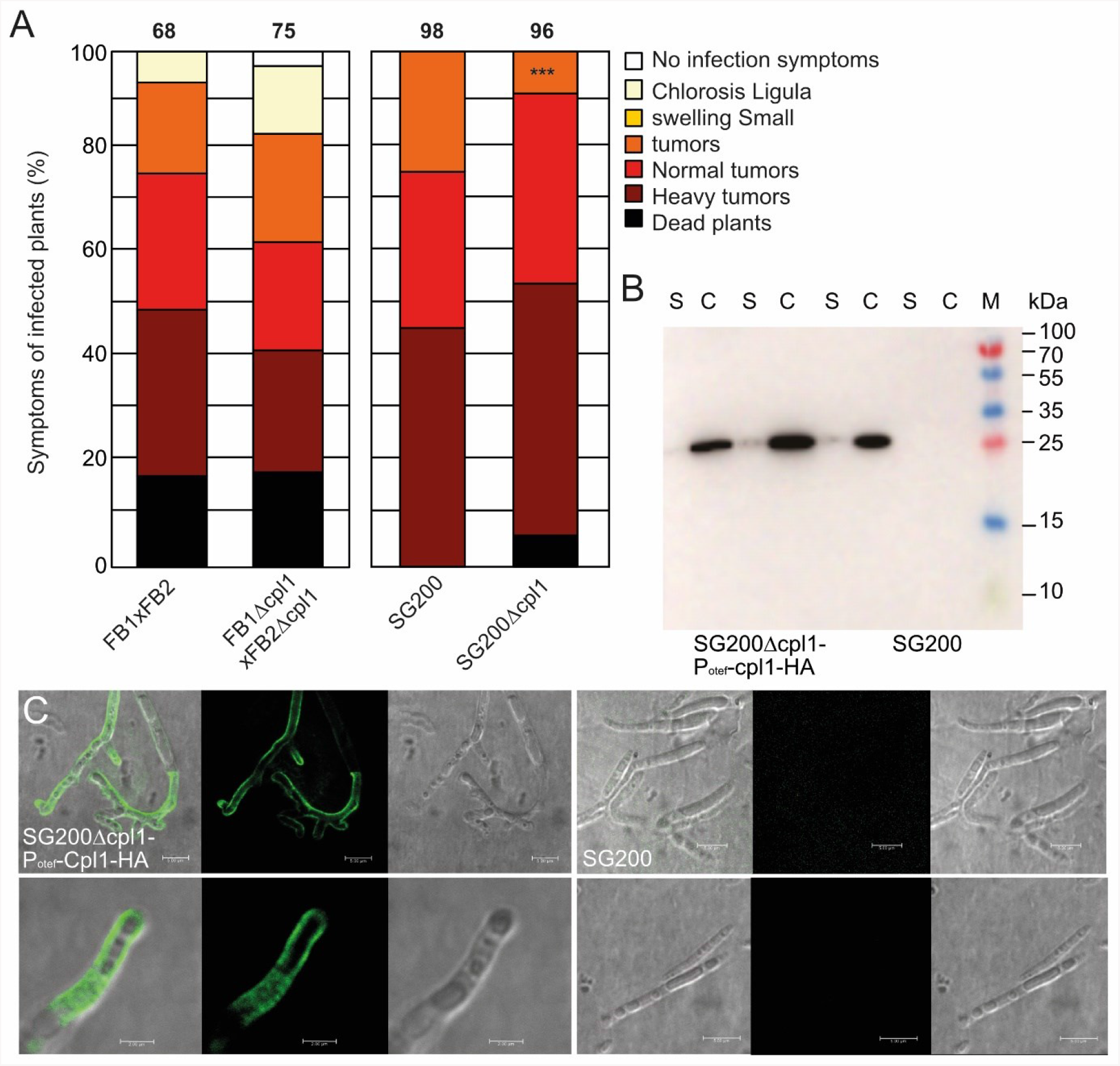
Cpl1 decorates the fungal cell wall of *U. maydis* hyphae. A. Maize infection assays using deletion strains in different genetic backgrounds of *U. maydis*. Cpl1 has been deleted from the genomes of FB1, FB2 and SG200 (from left to right). B. Western blot of SG200 strains overexpressing *cpl1-HA* shows that most Cpl1-HA stays attached to the cell wall and only a subfraction is found in the supernatant. S: Supernatant, C: Cell fraction C. Cpl1-HA is secreted and binds to the fungal cell wall of filaments grown on a parafilm surface. Confocal microscopy of strains immunostained with an anti-HA antibody and AF488-conjugated secondary antibody.

Although the available transcriptional data (Lanver et al., 2018) suggest that *cpl1* is not expressed under axenic conditions, we tested whether the growth of sporidial forms of these strains was affected under various stress conditions or if any morphological changes could be detected in sporidia. Here, we could neither observe differences in the cell morphology, nor differential responses towards the stress conditions tested (**Fig. S5A, B**). In conclusion, our data show that Cpl1 is dispensable for the growth of *U. maydis* in axenic culture and its deletion only mildly affects the virulence of *U. maydis*.

### Cpl1 localizes to the fungal cell wall and interacts with other cell-wall-associated proteins during infection

To consolidate our findings that Cpl1 binds to soluble chitin oligomers which suggests that it may localize to the fungal cell wall, we complemented the SG200 *cpl1* deletion strain by constitutively expressing *cpl1* fused to a C-terminal HA-tag. Strains overexpressing *cpl1-HA* were grown in liquid culture, harvested and the culture supernatant was precipitated using trichloroacetic acid, while the cells were subsequently lysed. Only faint amounts of Cpl1-HA were detected in the concentrated supernatant samples (**Fig. 4B**) but prominent amounts of Cpl1 could be detected in the cell samples (**Fig. 4B**). Again, overproduction of Cpl1-HA had no influence on the morphology of *U. maydis* sporidia or influenced the growth under different stress conditions (**Fig. S5**). To analyze if Cpl1-HA indeed resides in the cell wall, we stimulated strains with hydroxy-fatty acids and sprayed them on Parafilm to induce filamentation. The filaments were subjected to immunostaining using an anti-HA primary antibody and an Alexa Fluor 488 (AF488) conjugated secondary antibody. We detected evenly distributed fluorescence on long and branched filaments, but also shorter filaments showed a homogenous distribution of fluorescence (**Fig. 4C**). Interestingly, we did not detect any fluorescence in the areas surrounding the hyphae indicating that Cpl1-HA was tightly bound to the fungal cell wall. Taken together, our data support a role of Cpl1 in the fungal cell wall during maize infection.

Based on the localization of Cpl1-HA to *U. maydi*s filaments, we expected to potentially identify secreted maize proteins that might interact with Cpl1 *in planta*. This observation would be in line with Uvi2 from *U. hordei* interacting with a barley thaumatin in yeast two hybrid experiments (Ökmen et al., 2018). We therefore generated *cpl1-HA* complementation strains in the FB1 and FB2 deletion background to perform plant infection experiments followed by co-immunoprecipitation coupled mass spectrometry (**Fig. 5A**). We used an mCHERRY-HA under control of the promoter and fused to the signal peptide of Cmu1 as control. Both proteins could be purified from leaves of infected maize seedlings 3 days post infection and detected by western blotting (**Fig. 5B**).

**Figure 5.**
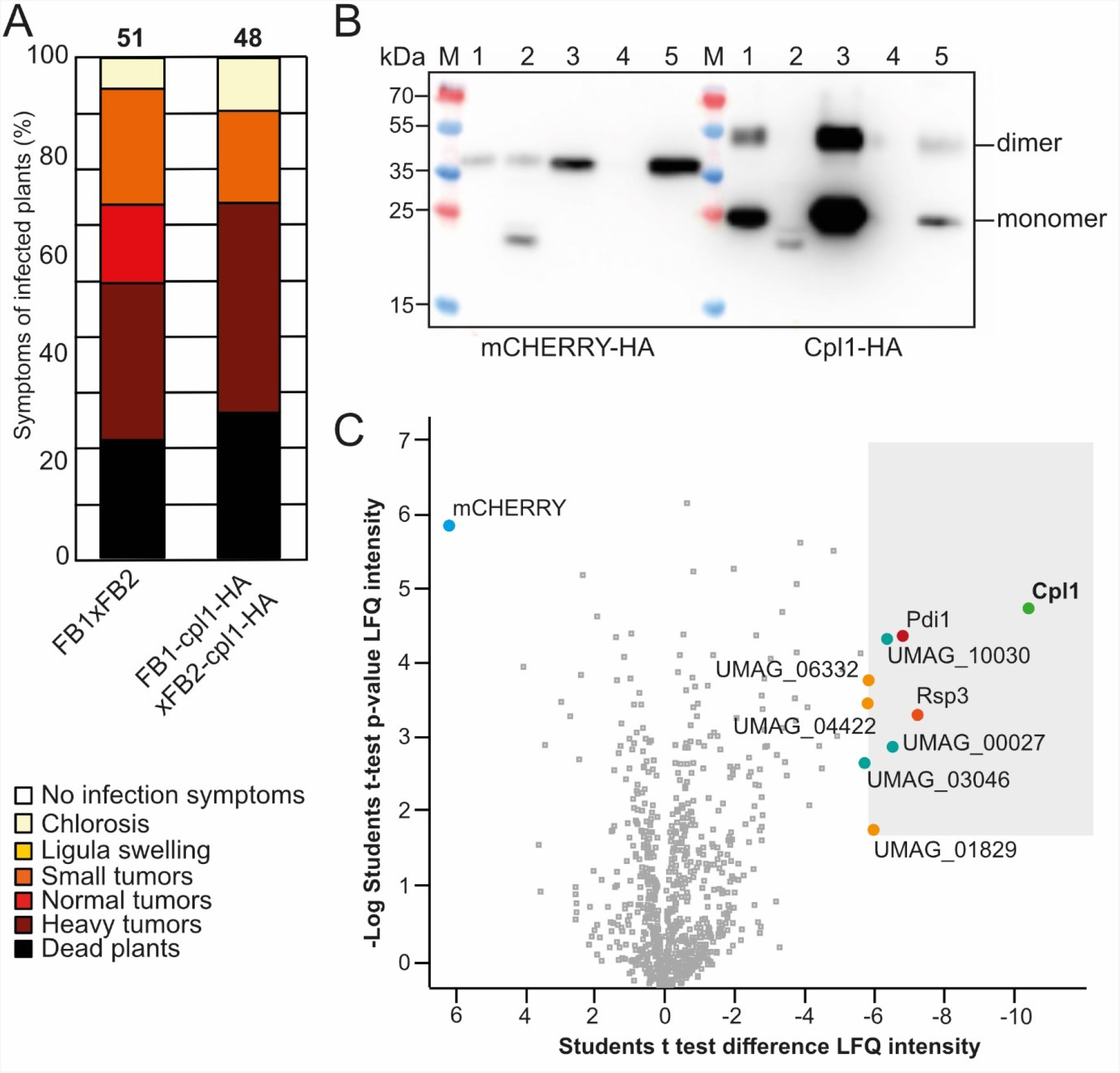
Cpl1 enriches cell wall degrading and cell wall decorating effectors during infection. A. Maize infection assay with the indicated *U. maydis* B. Western blot analysis of Cpl1-HA derived from infected maize leaves 3 days post infection. mCherry-HA fused to the signal peptide of Cmu1 under control of the *cmu1* promoter was used as a control. (1) Whole lysate of infected maize leaves, (2) insoluble fraction, (3) cleared lysate, (4) wash fraction and (5) Elution fraction. C. Volcano plot depicting the results of the third of the three Co-IP experiments conducted with FB1 and FB2 containing Cpl1-HA. Plant material infected with *U. maydis* containing mCherry-HA in the *ip*-locus under the promoter of *cmu1* was used as the control sample (see materials & methods). Shown are normalized label-free quantification (LFQ) intensities as either the p-value of the -Log Student’s t-test (y-axis) or the Student’s t-test difference. Data was acquired by liquid chromatography-mass spectrometry combined with the detection and quantification of peptide intensities (see materials & methods). A high -Log Student’s t-test p-value (y-axis) indicates a large number of peptides found, and a positive Student’s t-test difference (x-axis) means peptide specificity towards the bait sample containing Cpl1-HA (red and bold). All proteins enriched explicitly in the bait sample of all three Co-IP experiments are marked red. Proteins only significantly enriched in one or two of the three experiments are labeled black. For the control sample, only mCherry is marked (blue and bold).

To our surprise, eight *U. maydis* proteins were found to be significantly and repeatedly enriched in all three replicates based on their -Log (p-value) of the LFQ intensity, and their LFQ intensity difference. (**Fig. 5C, Tab. S5**). These proteins are UMAG_10030 (uncharacterized), UMAG_10156 (Pdi1, protein disulfide isomerase (Marín-Menguiano et al., 2019)) UMAG_06332 (Egl1, Endoglucanase 1 (Schauwecker et al., 1995)), UMAG_04422 (endo-1,4-β-xylanase (Moreno-Sánchez et al., 2021)), UMAG_03274 (Rsp3, cell wall-bound, protects against *Zm*AFP1/2 (Ma et al., 2018)), UMAG_00027 (uncharacterized), UMAG_03046 (uncharacterized), and UMAG_01829 (Afu1, putative non-reducing end α-L-arabinofuranosidase). Notably, all of them have a N-terminal signal peptide (SignalP 5.0), are rich in cysteine residues (average of 9.5 per protein, UniProt), and have no predicted transmembrane helices (THMHH 2.0; DTU Health Tech). These results suggest that Cpl1 localizes in the fungal cell wall in proximity or involved in a direct interaction with eight effector proteins during plant infection.

## Discussion

### Cpl1 – a novel type of cerato-platanin?

Plant colonization by pathogenic smut fungi is a complex multi-step process involving the secretion of a plethora of different virulence factors that are often species-specific and well-adapted to the respective host (Lanver et al., 2017; Zuo et al., 2019). In this work, we deliver a structural and biochemical characterization of a novel core effector protein from *U. maydis* that we termed Cpl1 due to the high structural similarity with proteins of the cerato-platanin (CP) family. CP proteins represent a group of expansin-related proteins found exclusively in filamentous fungi and are highly expressed during both filamentous growth on culture plates and on the surface of host plants (Baccelli, 2015; Luti et al., 2019; Narvaez-Barragan et al., 2020). These small, proteins typically have a central DPBB, four conserved cysteine residues and either reside in the fungal cell wall tightly bound to chitin or are secreted into the apoplast (Luti et al., 2019; Pazzagli et al., 2014). Although their precise role has not been clarified to date, they have been studied during plant colonization in several relevant plant pathogens, including *Botrytis cinerea, Magnaporthae grisea, Verticillium dahliae*, or *Fusarium graminearum* (Narvaez-Barragan et al., 2020). Interestingly, in some of these species, deletion of CP’s attenuated virulence while they were dispensable in others (Narvaez-Barragan et al., 2020). Notably, CP proteins have only been found in filamentous fungi and the fungi mentioned are prominent for their broad host range and aggressive hemibiotrophic or necrotrophic plant colonization behavior (Luti et al., 2019). Intriguingly, CP proteins are also linked to plant immunity, with many of them eliciting a defense response that triggers a hypersensitivity response, subsequently resulting in cell death and were thus regarded as PAMPs or MAMPs (Li et al., 2019; Luti et al., 2019; Narvaez-Barragan et al., 2020; Pazzagli et al., 2014). Our structure of Cpl1 now suggests that the CP-like fold as a distinguishing criterion of this protein family might be wider distributed among fungal species as anticipated so far.

### Cpl1 and Uvi2 might have divergent functions during plant infection

While Cpl1 shares the central DPBB and the chitin-binding properties with other CP-like proteins, there are some prominent differences. Chitin binding in CP’s is achieved through a flat and shallow surface groove that is conserved among this class of proteins (Chen et al., 2013). In contrast, this binding site is not conserved in Cpl1, where chitin oligomers bind in two grooves located within the dimer interface (**Figs. 2C, 3**). While CP’s form relatively compact particles, Cpl1 instead offers four loop-regions on both monomers (LR1-LR4, **Fig. 1B**) that result in the overall expanded appearance of Cpl1. We reason that these loop-regions might provide a modular binding interface for interaction partners like members of the plant protein family of kiwellins (Altegoer et al., 2020; Bange & Altegoer, 2019). Indeed, in a previous study Ökmen and coworkers performed yeast two-hybrid experiments and identified a barley thaumatin as an interaction partner of Uvi2, the Cpl1 homolog from *U. hordei* (Ökmen et al., 2018). Interestingly, deletion of *uvi2* led to a substantial decrease in fungal biomass during infection of barley by *U. hordei* (Ökmen et al., 2018), which suggests functional differences between Cpl1 and Uvi2. To investigate these differences in more detail which might allow explaining the different phenotypes in the two smut systems, we solved the crystal structure of Uvi2 (**Tab. S1; Fig. S6A**). The two structures superposed very well with an r.m.s.d of 0.548 covering all Cα-atoms (**Fig. S6A**). The central DPBB and both grooves on the upper and lower side of the protein were conserved on both sequence and structure levels. The highest structural deviation could be observed in the loop regions with the lowest sequence similarity (**Figs. S6A, B**). This further suggests that sequence variation in these loop-regions might facilitate the binding of specific interaction partners. In our Co-IP experiments Cpl1 co-enriched several fungal effectors during maize infection however, we could not detect peptides corresponding to maize thaumatin proteins (**Fig. 5C**).

A more detailed inspection of the proteins identified by Co-IP revealed the presence of three glycoside hydrolases, namely UMAG_04422 (Xyn1, β-Xylanase), UMAG_01829 (Afu1, Arabino-furanosidase), and UMAG_06332 (Egl1, Endoglucanase) (Moreno-Sánchez et al., 2021; Schauwecker et al., 1995). These cell wall degrading enzymes play important roles in cell wall remodeling during hyphal growth and plant cell penetration. Especially in the early stages of infection, removal of L-arabinose groups from arabinoxylans incorporated into the plant cell wall as a defense mechanism has shown to be an important step to increase the accessibility of xylan and penetrate the plant cell wall (de Vries et al., 2000; Doehlemann et al., 2008). The remodeling of fungal and plant cell walls needs to be well-orchestrated to prevent a host defense signaling. Taking the modular architecture of Cpl1 into account, a possible role might be the spatial organization of enzyme activity of these glycoside hydrolase enzymes during plant infection.

In contrast to necrotrophs and hemibiotrophs, biotrophic pathogens such as *U. maydis* have a reduced repertoire of cell wall degrading enzymes and might use these enzymes in a more fine-tuned manner (Spanu et al., 2010). An unrestrained enzymatic activity of these effectors might explain the slight hypervirulence observed upon deletion of *cpl1* (**Fig. 4A**). Furthermore, Cpl1 also enriched the disulfide isomerase Pdi1 which was shown to be important for quality control of cysteine-rich effectors during infection (Marín-Menguiano et al., 2019). Pdi1 resides in the endoplasmic reticulum of *U. maydis* sporidia but a significant amount is also localized to the cell wall (Marín-Menguiano et al., 2019). As our Co-IP experiments do not discriminate between Cpl1 that travels on the secretory pathway and mature cell wall-bound molecules, we can only speculate where a potential interaction with Pdi1 occurs. In addition, we could identify several peptides corresponding to Rsp3 which is an important cell-wall decorating protein shielding infectious hyphae from the activity of two mannose-binding proteins (Ma et al., 2018). Fungi have evolved several lines of defense to both protect their cell wall against attacking plant enzymes and prevent the generation of MAMPs that would elicit a plant immune response (Tanaka & Kahmann, 2021). Together with Rsp3, Cpl1 might also have a function in either shielding fungal hyphae or scavenging chitin fragments. As further effectors of yet known function were identified in our Co-IP experiments, more information on their function and how they connect to Cpl1 will likely help to understand the precise role of Cpl1 for virulence of *U. maydis*.

The interplay between effector proteins produced by smut fungi during infection is still poorly understood. Often, single deletion mutants are readily compensated by redundancy that masks the underlying function of the individual proteins. In humans, at least an estimated 22 % of all proteins are part of complexes (Giurgiu et al., 2019), and there is evidence that some effector proteins of *U. maydis* might be part of larger complexes (Alcantara et al., 2019). Slight variations in the amino acid sequence might not only alter the interactome but also allow for e.g. immune escape as shown for a stem rust effector recently (Ortiz et al., 2022). Our findings on Cpl1 with Uvi2 shed light on the role of a conserved effector with potentially diverging functions during plant infection by *U. maydis* and *U. hordei* due to subtle amino acid variations.

## Supporting information

Supplemental_Dataset1-HDX

Supplemental_Dataset2-MScounts

Supplemental_Dataset3-LFQ-data-matrix

## Acknowledgements

We thank Regine Kahmann for her continuous support throughout the work on this manuscript and her group for helpful discussions. We also thank Mariana Schuster for expert help with the CRISPR-Cas9 system, Xiaowei Han for assistance with strain construction and plant infection experiments. We thank Vera Göhre and Michael Feldbrügge for critical reading and fruitful discussions on the manuscript. We thank the EMBL Hamburg at the PETRA III storage ring (DESY, Hamburg, Germany) for support. G.B. thanks the European Research Council (ERC) for support through the project “KIWIsome” (Grant agreement number: 101019765).

## Author contributions

F.A. and G.B. conceived of the project and designed the study. F.A. and P.W. wrote the paper with input from all other authors. P.W, F.D., W.S., T.G. and R.M. performed experiments. P.W., F.D., S.A.F, W.S., T.G. and F.A. analyzed data. G.B. contributed funding and resources.

## Data availability

Coordinates and structure factors have been deposited within the protein data bank (PDB) under accession codes: 8A14 and 8A4O. The authors declare that all other data supporting the findings of this study are available within the article and its supplementary information files.

