## Supplementary Information for "Structural and functional analysis of the cerato-platanin-like effector protein Cpl1 suggests diverging functions in smut fungi"

**Table S1.** Crystallographic data collection and refinement statistics

| | Cpl1 (8A14) | Uvi2 (8A4O) |
|---|---|---|
| **Data collection** | | |
| Space group | $P2_12_12_1$ | $P6_1\ 2\ 2$ |
| Cell dimensions | | |
| *a*, *b*, *c* (Å) | 85.71 121.92 180.9 | 69.05 69.05 232.59 |
| α, β, γ (°) | 90 90 90 | 90 90 120 |
| Resolution (Å) | 49.68 - 1.797 (1.861 - 1.797) | 36.72 -1.35 (1.401 - 1.353) |
| $R_{merge}$ | 0.1165 (1.929) | 0.1214 (4.728) |
| $I / \sigma I$ | 16.03 (1.53) | 17.93 (0.51) |
| Completeness (%) | 98.97 (90.01) | 99.75 (98.28) |
| Redundancy | 13.5 (13.4) | 35.2 (21.9) |
| $CC_{1/2}$ | 0.999 (0.645) | 1 (0.416) |
| **Refinement** | | |
| Resolution (Å) | 49.68 - 1.797 | 36.72 – 1.35 |
| No. reflections | 174511 (15743) | 72482 (6967) |
| $R_{work}$ / $R_{free}$ | 0.17/0.19 | 0.16/0.18 |
| No. atoms | | |
| Protein | 9669 | 1615 |
| Ligand/ion | - | - |
| Water | 1475 | 299 |
| *B*-factors | | |
| Protein | 31.71 | 24.84 |
| Ligand/ion | - | - |
| Water | 42.85 | 37.37 |
| Ramachandran (%) | | |
| favored | 97.84 | 96.86 |
| allowed | 2.07 | 3.14 |
| outliers | 0.00 | 0.00 |
| R.m.s. deviations | | |
| Bond lengths (Å) | 0.011 | 0.025 |
| Bond angles (°) | 1.64 | 1.76 |

*Values in parentheses are for highest-resolution shell.

**Table S2.** Plasmids used in this study.

| ***Plasmid*** | ***Usage*** | ***Citation*** |
|---|---|---|
| *pET24d* | Protein overexpression with N-terminal 6His-tag | (Novagen) |
| *pEMGB1* | Protein overexpression with N-terminal GB1-tag | (Zhou & Wagner, 2010) |
| *pMS73* | CRISPR/Cas9 vector for mutliplexed genome editing in *Ustilago maydis*. Addgene #110629 | (Schuster et al., 2018) |
| *pFA010* | pMS73 with sgRNA to disrupt *UMAG_01820* for gene deletion | (This study) |
| *pFA013* | Plasmid contains *cpl1-HA* under control of the constitutive otef promoter to be introduced in the *ip* locus of *U. maydis* | (This study) |
| *pFA438* | Overexpression of Uvi2 with an N-terminal hexahistdine tag | (This study) |

| | | |
|---|---|---|
| *pFA451* | Overexpression of Cpl1 with N-terminal GB1-tag separated from the ORF by a TEV cleavage site | (This study) |
| *pFA576* | pMS73 with sgRNA to disrupt *UMAG_01820* for C-terminal tagging | (This study) |
| *pPW126* | Overexpression of Cpl1(mut1 - N154A-R158A) with N-terminal GB1-tag separated from the ORF by a TEV cleavage site | (This study) |
| *pPW127* | Overexpression of Cpl1(mut2 - D56A-E59A) with N-terminal GB1-tag separated from the ORF by a TEV cleavage site | (This study) |

**Table S3.** Primers used in this study.

| ***Number*** | ***Sequence*** | ***Description*** | ***Reference*** |
|---|---|---|---|
| *oMS74-Rv* | CGGCGTTCGACTCTT | Reverese primer to generate pFA010 and pFA576 | (Schuster et al., 2016) |
| *oFA363* | AGGAGGGTCTCCCATGGGCAGCATGGACATCACATTC | Forward Primer used to generate pFA438 | (This study) |
| *oFA365* | AGGAGGGTCTCCTCGAGTTAACCGATGGTGTTAATATC | Reverse Primer used to generate pFA438 | (This study) |
| *oFA383* | TCCTCGGTCTCCTCGAGTTAACCCACGGTGTTGATGTCGAG | Reverse Primer used to generate pFA451 | (This study) |
| *oFA504* | CAAAATTCCATTCTACAAC**G**GTCGAGGAAGAACCACTCGAGTTTTAGAGC | Forward primer to generate pFA576 | (This study) |
| *oFA505* | TCCGGTCGAGTGGTTCTTCCTCGACATCAACACCGTGGGTGGATCCtacccctacgacgtgcccgactatgccTAGTTGGATTTCTCCCTATTCAGCTTCTGCGAATCCTGAA | Donor DNA for generating *cpl1-HA* | (This study) |
| *oFA729* | AGGAGGGTCTCCCATGGGCGCCGTTGACATCACTTTTACCTCG | Forward Primer used to generate pFA451 | (This study) |
| *oFA730* | AGGAGTCATGAGCAAGTTTGAATTTGGTGCCCTTGTC | Forward primer to generate pFA013 | (This study) |
| *oFA731* | AGGAGgcggccgcTCAggcatagtcgggcacgtcgtaggggtaGGATCCACCCACGGTGTTGATGTCGAG | Reverse primer to generate pFA013 | (This study) |
| *oPW188* | AGGAGGGTCTCGCGCTGTACGACGCGCGCT | Forward Primer used to generate pPW126 | (This study) |
| *oPW189* | AGGAGGGTCTCCAGCGCGTTCATGTAGAGGTAAGGAACA | Reverse Primer used to generate pPW126 | (This study) |
| *oPW190* | AGGAGGGTCTCAAGGCGACTTACCCCGAGTGCAAGTGG | Forward Primer used to generate pPW127 | (This study) |
| *oPW191* | AGGAGGGTCTCCGCCTTGACGGCCTTGGCCGG | Reverse Primer used to generate pPW127 | (This study) |

| | | |
|---|---|---|
| *oFA420* | CAAAATTCCATTCTACAACGGCC AGGTTCATGCCAAACTCGTTTTA GAGC | Forward primer to generate pFA010 |
| *oFA421* | TCTCCCCTGTTCACATACTCTGCTCT CGATCCTAGTCAAGTTGGATTTCTC CCTATTCAGCTTCTGCGAATCCTGA ATTT | Donor DNA for generating *cpl1* deletion strains |

**Table S4.** Strains used in this study.

| ***Strain*** | ***Parental strain*** | ***Vector*** | ***Genotype*** | ***Citation*** |
|---|---|---|---|---|
| SG200 | - | - | *a1 mfa2 bW2 bE1* | (Kämper et al., 2006) |
| FB1 | - | - | *a1b1* | (Banuett & Herskowitz, 1989) |
| FB2 | - | - | *a2b2* | (Banuett & Herskowitz, 1989) |
| SG200mCherry-HA | SG200 | - | *a1 mfa2 bW2 bE1* $ip^R[P_{cmu1}:SP_{cmu1}:mcherry:biotag:HA]ip^S$ | (Lo Presti et al., 2017) |
| FA027 | SG200 | pFA10 | *a1 mfa2 bW2 bE1 Δcpl1* | (This study) |
| FA035 | FB1 | pFA10 | *Δcpl1* | (This study) |
| FA038 | FB2 | pFA10 | *Δcpl1* | (This study) |
| FA063 | FB1 | pFA576 | $P_{cpl1}$*:cpl1-HA* | (This study) |
| FA064 | FB2 | pFA576 | $P_{cpl1}$*:cpl1-HA* | (This study) |
| FA083 | SG200 | pFA14 | *a1 mfa2 bW2 bE1* $ip^R$ [$P_{otef}$:*cpl1-HA*] $ip^S$ | (This study) |

**Table S5. All eight significantly and repeatedly enriched proteins with their LFQ values.** The table below lists the p-value of the -log10 Student's t-test and the Student's t-test difference of the measured and normalized label-free quantitation (LFQ) intensities for each of the three Co-IP experiments.

| **protein** | **-Log (p-value)** Co-IP I | Co-IP II | Co-IP III | **standard deviation** | **difference** Co-IP I | Co-IP II | Co-IP III | **standard deviation** |
|---|---|---|---|---|---|---|---|---|
| Egl1 | 3.76 | 4.83 | 4.44 | 0.44 | 4.38 | 5.50 | 5.12 | 0.47 |
| UMAG_00027 | 3.40 | 5.96 | 3.20 | 1.26 | 6.97 | 5.34 | 5.90 | 0.67 |
| Cpl1 | 6.04 | 8.04 | 5.03 | 1.25 | 12.66 | 9.08 | 9.45 | 1.61 |
| Afu1 | 3.28 | 2.35 | 2.12 | 0.50 | 6.75 | 5.11 | 5.46 | 0.71 |
| UMAG_03046 | 4.33 | 3.96 | 2.97 | 0.58 | 7.07 | 4.26 | 5.21 | 1.17 |
| Rsp3 | 6.89 | 3.68 | 3.59 | 1.53 | 6.57 | 5.93 | 6.58 | 0.30 |
| UMAG_04422 | 5.58 | 4.81 | 4.07 | 0.62 | 5.65 | 5.67 | 5.29 | 0.18 |
| UMAG_10030 | 4.57 | 4.53 | 4.60 | 0.03 | 6.14 | 5.81 | 5.79 | 0.16 |
| Pdi1 | 4.83 | 4.76 | 4.65 | 0.07 | 8.98 | 7.34 | 6.20 | 1.14 |

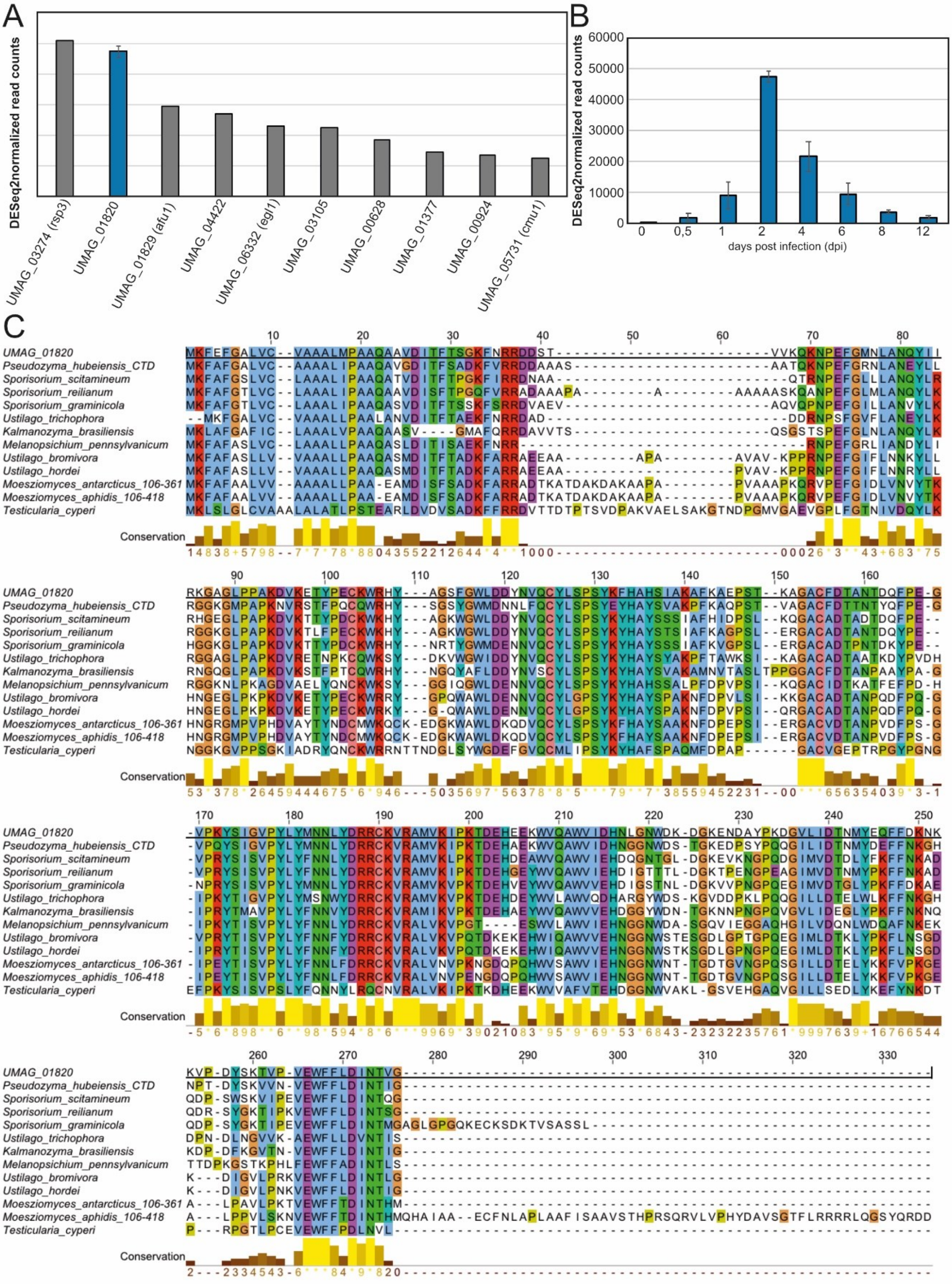

**Supplementary Figure 1. Multiple sequence alignment of Cpl1 homologs. A.** The ten most abundant transcripts at 2 days post infection. Data were taken from (Lanver et al., 2018). **B.** Expression profile of *cpl1* during plant infection. Shown are normalized read counts taken from (Lanver et al., 2018). **C.** The aa sequences of all identified orthologs were aligned against the sequence of Cpl1 to highlight similarities. Conservation scores and visualization were done using Jalview (Waterhouse et al., 2009). Sequence alignment was done using CLUSTAL2.1 (Sievers et al., 2011). The aa coloring scheme is based on Clustal. For better visualization, only the C-terminal region of *P. hubeiensis* SY62 was used, and for *M. antarcticus* and *M. aphidis* aa 106 – 361 and 106 – 418, respectively.

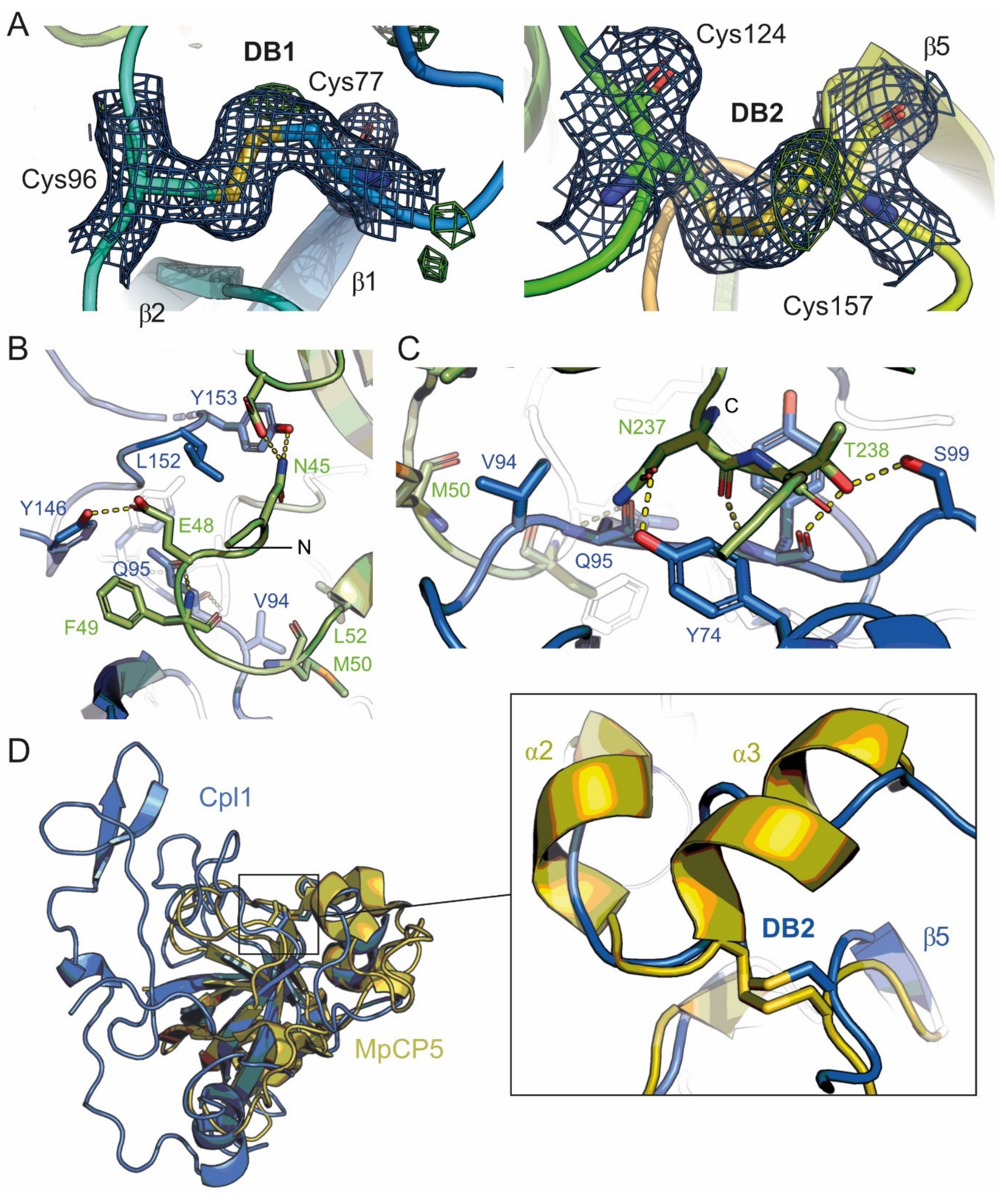

**Supplementary Figure 2. Cpl1 has conserved disulfide bonds. A.** Closeup of the two disulfide bonds (DB) of Cpl1 depicted as sticks with the respective $2F_{obs}$-$F_{calc}$ electron density maps contoured at 2.0 σ. **B.** and **C.** Detailed view of the dimer interface between the two molecules of Cpl1. Dashed lines indicated polar interactions. Green and blue color corresponds to the two monomer molecules. **D.** Superposition of Cpl1 (blue) and MpCP5 (PDB-Code: 3SUM; yellow) shows that DB2 of Cpl1 superposes well with one of the DBs from MpCP5.

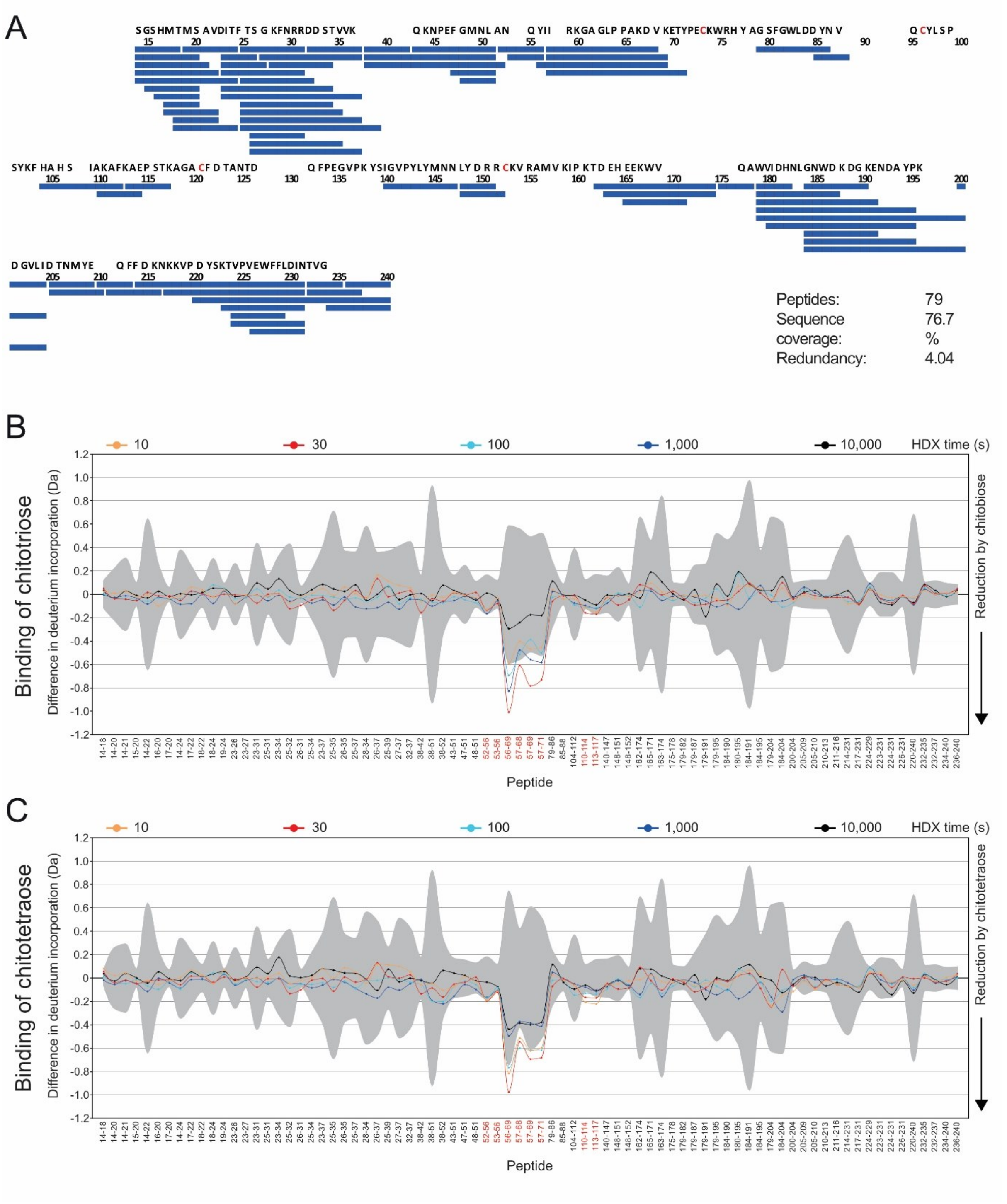

**Supplementary Figure 3. Binding of chitobiose and chitotetraose to Cpl1 by HDX-MS. A.** Peptides of Cpl1 that were analyzed for their deuterium incorporation are indicated as blue bars and plotted on the amino acid sequence of Cpl1. The disulfide bridge-forming cysteine residues are colored in red. **B.** and **C.** The difference in deuterium incorporation between chitobiose-bound (**B**) and chitotetraose-bound (**C**) Cpl1 versus apo-Cpl1 is displayed per peptide. Grey color denotes a 3$\sigma$ multiplier error band (n=3) based on the SD's of each peptide. Negative values of the difference represent reduced deuterium incorporation of Cpl1 in presence of chitobiose or chitotetraose. The plots were generated with MEMHDX (Hourdel et al., 2016).

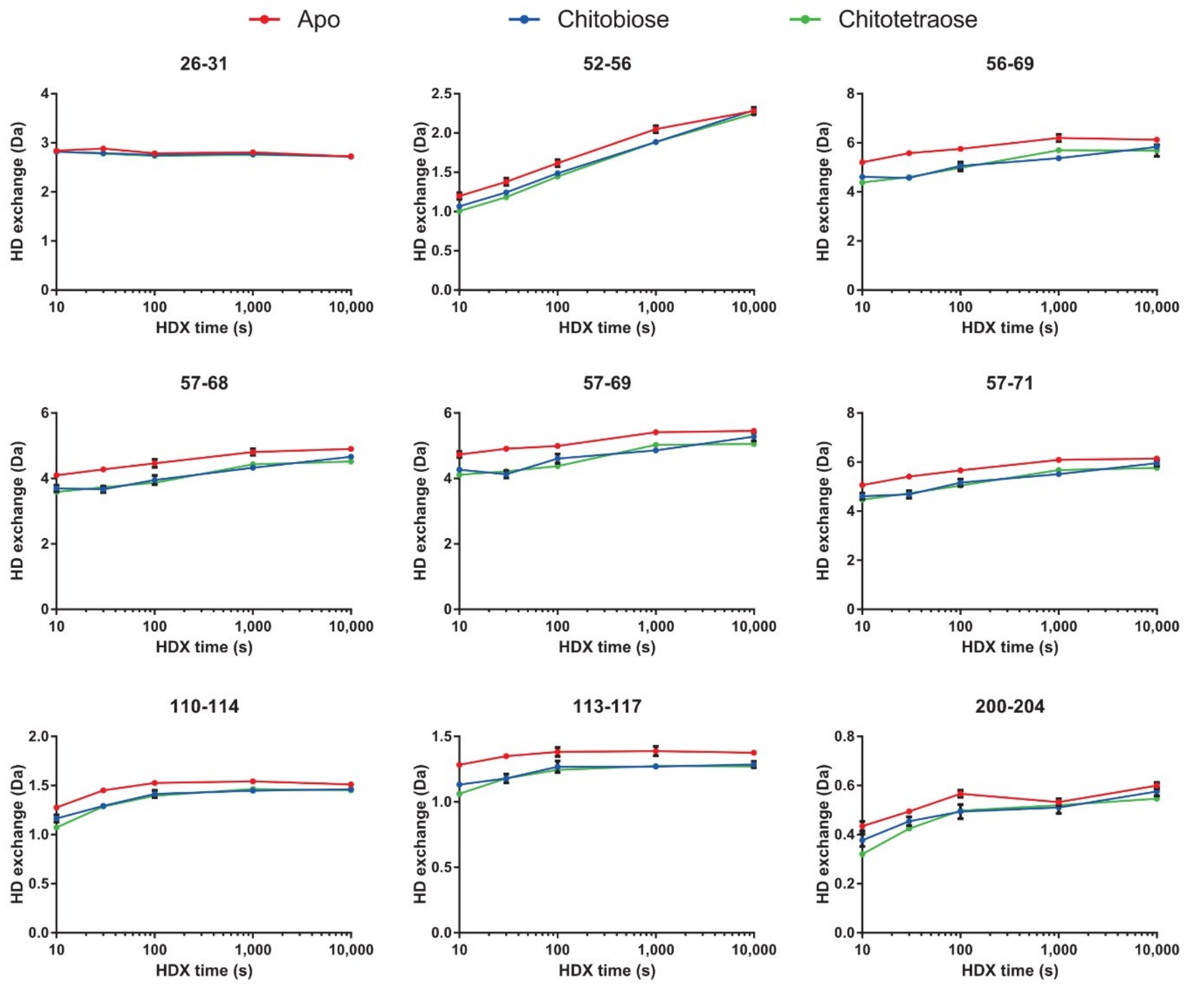

**Supplementary Figure 4.** Binding of chitobiose and chitotetraose to Cpl1 by HDX-MS. A. Deuterium incorporation of representative Cpl1 peptides in the absence of ligand (red) or in the presence of chitobiose (blue) or chitotetraose (green). Data represent the mean ± SD of three technical replicates (individual HDX reactions) for each time-point.

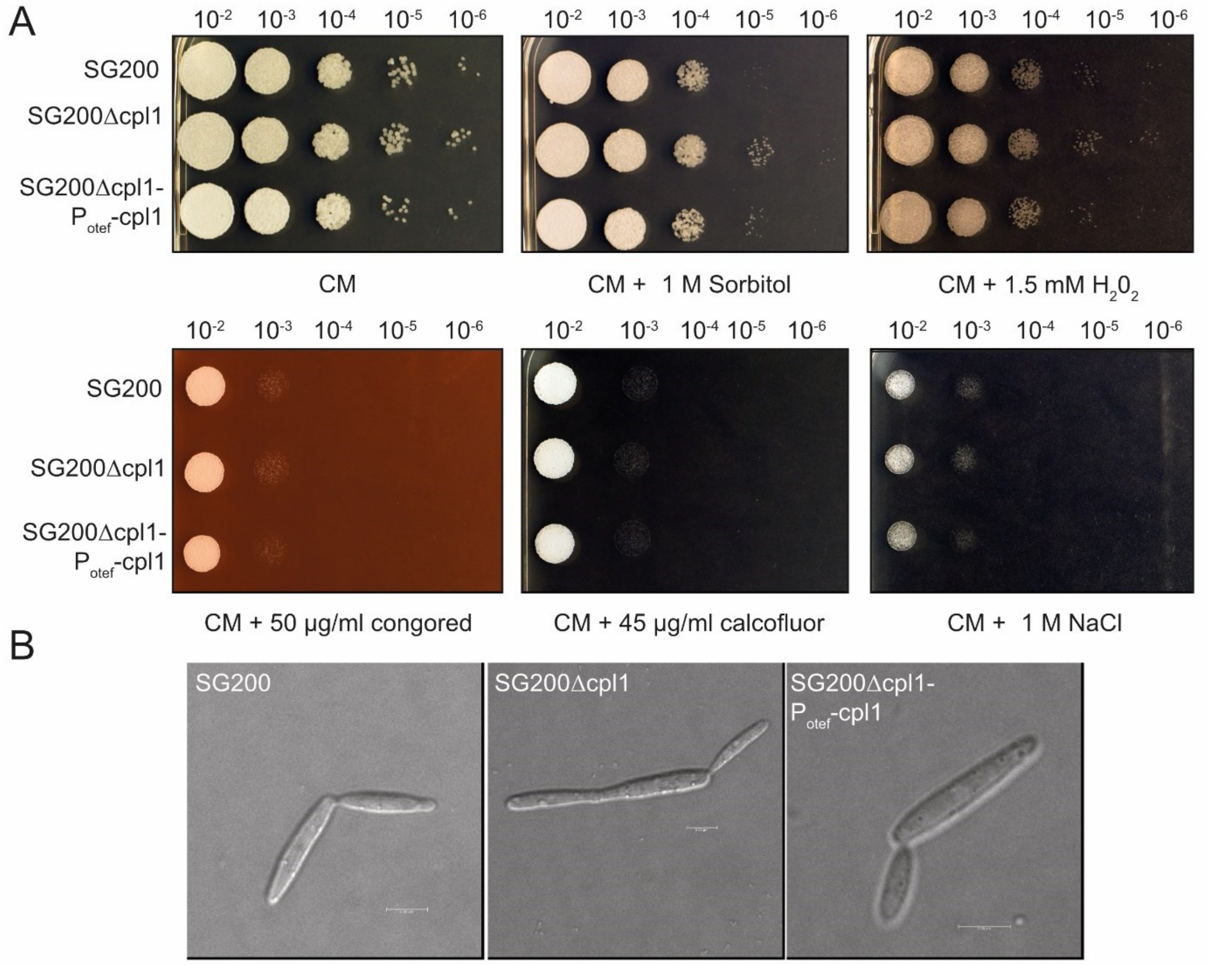

**Supplementary Figure 5. Stress assays. A.** Stress assays of SG200, SG200Δcpl1, SG200Δcpl1-$P_{otef}$-cpl1 were performed on CM plates containing sorbitol, $H_2O_2$, congored, calcofluor white and NaCl. **B.** Microscopy of SG200 and the respective *cpl1* deletion strains and strains complemented with *cpl1* under control of the constitutive *otef* promoter.

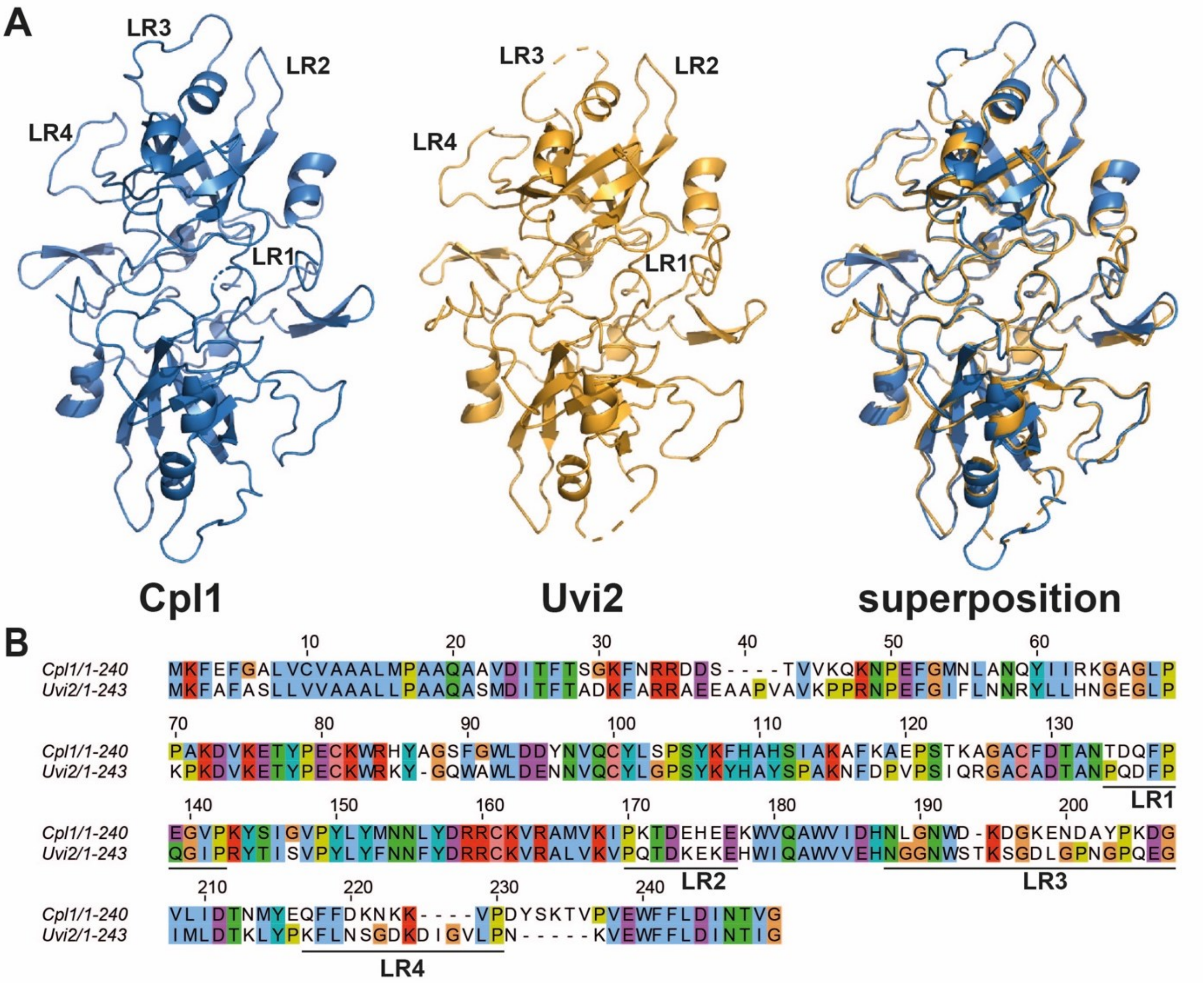

**Supplementary Figure 6. Structural comparison of Cpl1 and Uvi2. A.** Superposition of Cpl1 and Uvi2 structures shows that the overall architecture is conserved, and small structural deviations relate to the loop regions. **B.** Amino acid sequence alignment of Cpl1 and Uvi2. The loop regions have the highest sequence deviations.
